# MAPPER: A new image analysis pipeline unmasks differential regulation of *Drosophila* wing features

**DOI:** 10.1101/2020.12.16.422888

**Authors:** Nilay Kumar, Francisco Huizar, Trent Robinett, Keity J. Farfán-Pira, Dharsan Soundarrajan, Maria Unger, Pavel Brodskiy, Marcos Nahmad, Jeremiah J. Zartman

**Affiliations:** Department of Chemical and Biomolecular Engineering, University of Notre Dame, Notre Dame, IN 46556, USA; Department of Physiology, Biophysics, and Neurosciences, Center for Research and Advanced Studies of the National Polytechnical Institute (Cinvestav), Mexico City, Mexico 07360.

## Abstract

Phenomics requires quantification of large volumes of image data, necessitating high throughput image processing approaches. Existing image processing pipelines for *Drosophila* wings, a powerful model for studying morphogenesis, are limited in speed, versatility, and precision. To overcome these limitations, we developed MAPPER, a fully-automated machine learning-based pipeline that quantifies high dimensional phenotypic signatures, with each dimension representing a unique morphological feature. MAPPER magnifies the power of *Drosophila* genetics by rapidly identifying subtle phenotypic differences in sample populations. To demonstrate its widespread utility, we used MAPPER to reveal new insights connecting patterning and growth across *Drosophila* genotypes and species. The morphological features extracted using MAPPER identified the presence of a uniform scaling of proximal-distal axis length across four different species of *Drosophila*. Observation of morphological features extracted by MAPPER from *Drosophila* wings by modulating insulin signaling pathway activity revealed the presence of a scaling gradient across the anterior-posterior axis. Additionally, batch processing of samples with MAPPER revealed a key function for the mechanosensitive calcium channel, Piezo, in regulating bilateral symmetry and robust organ growth. MAPPER is an open source tool for rapid analysis of large volumes of imaging data. Overall, MAPPER provides new capabilities to rigorously and systematically identify genotype-to-phenotype relationships in an automated, high throughput fashion.

**Graphical abstract:** 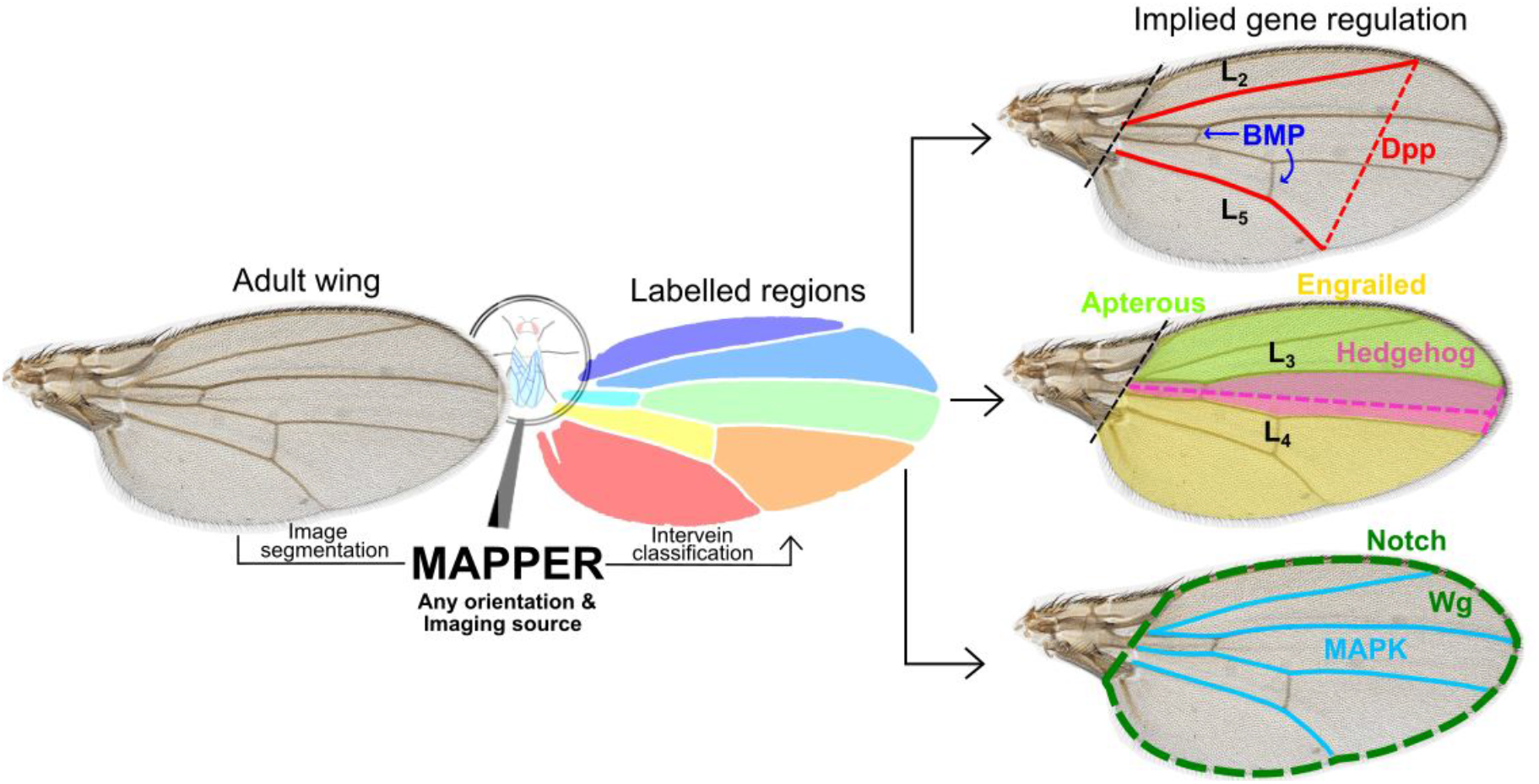

## Introduction

### The challenge of phenomics in multicellular organs

The architectural maxim of L. Sullivan “form follows function” is rigorously observed in many biological structures where shape is a key determinant of function (Sullivan 1896). Mapping the functional relationships between genotypes and phenotypes involves translating phenotypic data typically available as an image into a high dimensional space that describes key morphometric features. Quantification and subsequent comparison of morphometric features is crucial for identifying and explaining gene conditions responsible for the phenotype. Advances in imaging and machine learning (ML) empower the application of phenomics in a high throughput fashion due to the ease of identification of patterns in features (Houle et al. 2017).

The *Drosophila* wing has an excellent track record for genetic screening studies and is ideal for phenomic studies to uncover conserved biological processes relevant to human development and diseases. The *Drosophila* wing has successfully identified genes crucial for organ development and relevant to human health (Buchmann, Alber, and Zartman 2014; Narciso and Zartman 2018; Restrepo, Zartman, and Basler 2014; Strigini and Cohen 1999; Bier 2005; K. Kim et al. 2020; Brock, Seto, and Smith-Bolton 2017). Further, the developing wing imaginal disc has often been used for studying growth, development, and tissue regeneration (Jaszczak and Halme 2016; Smith-Bolton et al. 2009; Hariharan and Serras 2017). The *Drosophila* wing blade includes five longitudinal veins, two cross veins, intervein trichomes, and hairs along the surface and edge of the wing. These visual features provide a flat readout of conserved signaling pathway activity (Figure 1A, Figure S1) (Bier 2005). Wing development is a systems-level process that requires coordinated regulation of cellular processes such as proliferation, differentiation, and morphogenesis (Huizar et al. 2020; Restrepo, Zartman, and Basler 2014; Neto-Silva, Wells, and Johnston 2009; Diaz de la Loza and Thompson 2017; De Celis 2003; O’Connor et al. 2006). The final shape and size of the adult wing depends on the integration of both intrinsic genetic regulatory network and extrinsic environmental cues such as temperature, nutrition, and hormones (Johnston and Gallant 2002; Parker and Struhl 2020).

**Figure 1.**
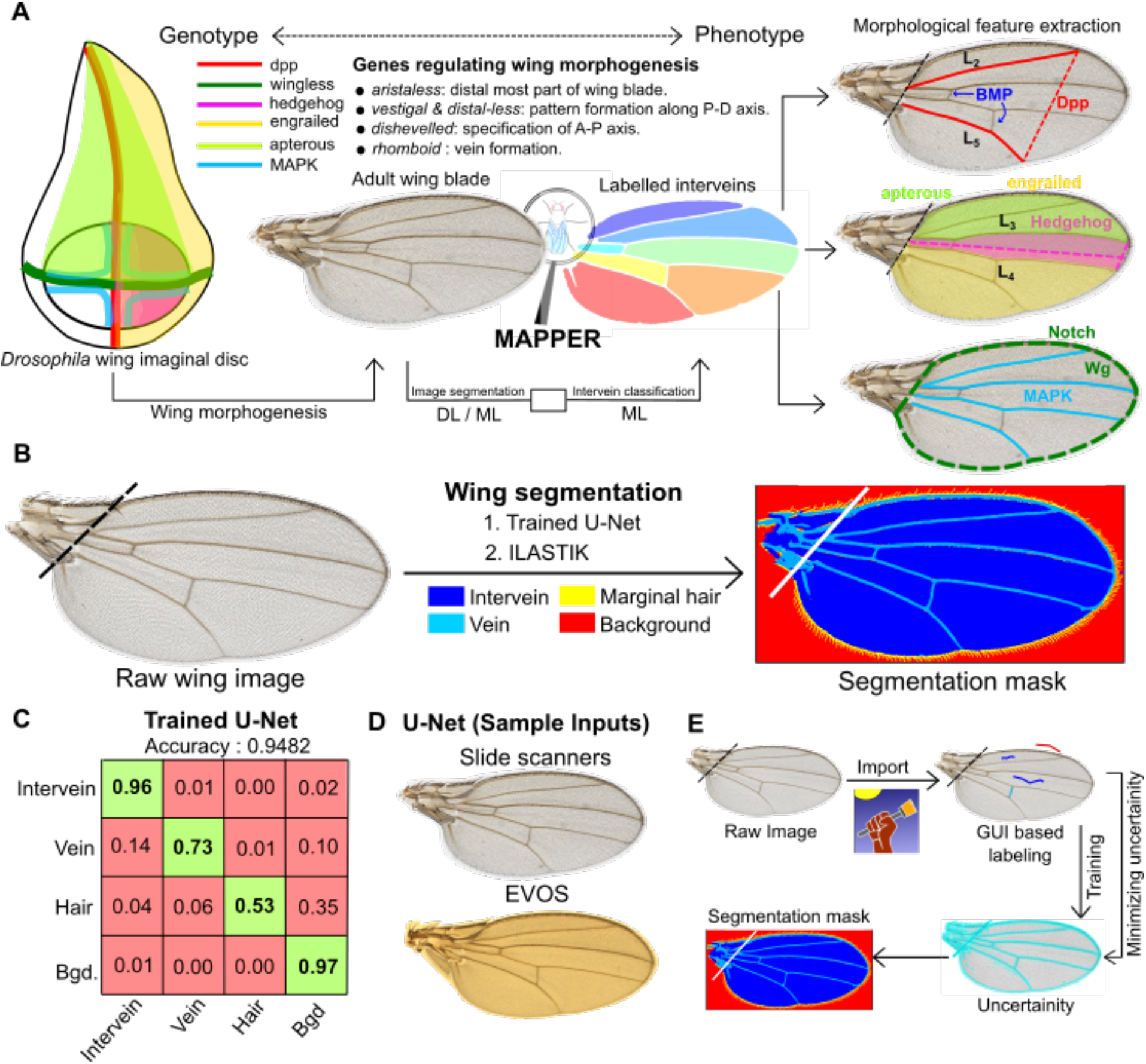
MAPPER automates segmentation of wings. **(A)** Overview of MAPPER where a *Drosophila* wing imaginal disc is regulated by various genes (Campbell, Weaver, and Tomlinson 1993; Boutros and Mlodzik 1999; Williams, Bell, and Carroll 1991; Gomez-Skarmeta et al. 1996), and becomes an adult wing blade through morphogenesis. The coupled image segmentation and intervein classification processes of MAPPER enables morphological feature extraction. **(B)** Image segmentation of adult *Drosophila* wings (left) is carried out for identification and labelling of different regions of interest (right) within the adult wing blade. Two methods of training are available: 1. The Trained U-Net or 2. ILASTIK. **(C)** MAPPER utilizes the U-Net architecture, which consists of convolutional layers for feature extraction, followed by deconvolution layers to achieve pixel- level prediction. Confusion matrix showing pixel classification accuracy for a U-net trained to identify different regions of interest. The numbers in the boxes represent the prediction accuracy for classification of a pixel into a class represented in the vertical labels against the true class in the horizontal labels. **(D)** Sample wings from different imaging sources that can be processed by a trained U-net model. **(E)** Schematic describing methodology followed by ILASTIK, an open source pixel classifier, for the purpose of pixel classification.

The success of the *Drosophila* wing as a model system for genotype-phenotype studies is due to its balance between structural simplicity and functional complexity. Subtle changes in the shape and size of the wing provide insight into conserved signaling mechanisms that occur during wing development (Gibson and Dworkin 2004; Kawecki et al. 2012; Matamoro-Vidal et al. 2018). However, most of the phenotypic studies carried out using *Drosophila* wings as a model system produce a large volume of image data. Such data traditionally has been analyzed manually or aided by semi-automated pipelines. Manually quantifying phenotypic features is extremely time consuming for large data sets produced in phenomic studies. This makes manual extraction of key morphometric features, such as wing size, shape, trichome density, and trichome distribution, impractical over the large sample sizes required to obtain reproducible results. Previous efforts have developed algorithms to perform high throughput analysis of these features (Dobens and Dobens 2013; Houle et al. 2003). However, they are still limited by a lack of computational speed, accuracy, and versatility in the quantification of morphometric traits and only extract a limited number of morphometric traits.

### The solution presented in MAPPER and why it’s transformative

To overcome the limitations of manual and semi-automated platforms, we developed the Multicellular Analysis Platform for Engineering Research (MAPPER), a fully-automated pipeline for insect wing segmentation and morphometric feature extraction. MAPPER is composed of two distinct modules that operate sequentially. First, the pipeline employs a deep learning (DL)-based image segmentation platform, where we trained a convolutional neural network, U-Net, for automated identification of different components of the wing. The trained DL model generates segmentation masks that define different regions of the wing, at a much faster rate compared to conventional image segmentation algorithms like active contours or image thresholding. Further, the model can also be re-trained with new images, easily making it more generalizable for datasets belonging to different imaging sources, thereby allowing versatility across various research labs. Alternatively, users can also use ILASTIK, a ML-based pixel classifier for the same task. Following the image segmentation pipeline, is a k-nearest neighbor (KNN)-based machine learning classifier (Cover and Hart 1967) that classifies and labels each intervein region. This facilitates high throughput feature extraction in an organized fashion. Identification and labelling of interveins allows MAPPER to extract hundreds of geometrical features in a systematic and high throughput manner. Together, these pipelines allow MAPPER to accurately and swiftly extract large amounts of phenotypic data from wing imaging datasets.

### Case studies demonstrating the capability, versatility, and implementation of MAPPER

To benchmark MAPPER with previous wing analysis packages, we used MAPPER to confirm the role of insulin receptors (InsR) in regulating *Drosophila* wings (Brogiolo et al. 2001). While insulin receptors are known to regulate size, MAPPER enabled discovery of the presence of a scaling pattern of intervein area along the anterior-posterior (AP) axis of the wing blade in InsR-mediated morphogenesis. MAPPER enabled a complete systematic analysis of how wing shape varies across four *Drosophila* species: *D. ananassae*, *D. melanogaster*, *D. simulans*, and *D. virilis.* MAPPER’s analysis revealed subtle differences, such as the scaling relationships between intervein regions, that would be very difficult to obtain from manual or semi-automated platforms. These observations shed light in the genetic regulatory processes underlying wing shape and size. Finally, we used MAPPER to identify local and global shape and patterning changes that result from perturbations to the mechanosensitive cation channel gene *Piezo*, providing new insight into the functional roles of mechanosensitive channels in organ size regulation. MAPPER is open source and available in the form of an interactive GUI, thus making the tool usable to users with no prior experience in programming.

## Design

Several image processing pipelines for phenotypic characterization of adult *Drosophila* wings have been proposed in the past which include open source tools like FIJIWings (Dobens and Dobens 2013) and WINGMACHINE (Houle et al. 2003). However, existing pipelines are insufficient as high throughput characterization tools, making it difficult to identify pattern-based variations within wing populations. Further, previous pipelines employ conventional image segmentation algorithms, such as thresholding, for the identification of different regions of interest within the wing. Such algorithms are sensitive to noise and also require tuning of algorithm-based parameters. Adding to this, some pipelines require the wing to be imaged in a specific orientation for estimation of landmark features, further increasing the processing time involved in reorienting images. Recent work has proposed rotation-free estimation of landmark coordinates (Alba et al. 2020). However, this approach does not quantify local variations in patterning of the peripheral shape of the wing. The existing pipelines have not fully utilized the recent advancements made in the field of statistical learning, which is notably advancing biomedical image analysis (Klang 2018; M. Kim et al. 2019). MAPPER utilizes these algorithmic advances for robust high throughput analysis of *Drosophila* wings. Here, we re-calibrated a DL-based network, U-Net (Ronneberger, Fischer, and Brox 2015), to separate intervein and veins from the imaging background. A ML-based classifier then labels each intervein region based on their shape. Intervein classification by U-Net is followed by extraction of features that describe the shape of the wing using Elliptic Fourier Descriptors (EFDs) (Kuhl and Giardina 1982). EFDs are defined to measure local and global changes in the overall shape of wing. The labeling of interveins provides an orientation-free classification of veins, which is followed by estimation of landmark features and anatomical axes, such as the proximal-distal (PD) axis and the anterior-posterior (AP) axis. MAPPER is designed for use by a broad user base of traditional biologists and bioengineers. MAPPER is available in the form of a MATLAB-based GUI for both individual and batch analysis of wings. The design of MAPPER also allows users with preliminary knowledge of MATLAB to integrate their custom functions estimating any new desired geometric feature. Details about the design, use and application of MAPPER are provided below (SI Section S1 - S4).

## Results

### MAPPER uses statistical learning techniques to automate identification of *Drosophila* wing features

Automation of any quantitative feature extraction pipeline depends primarily on the accuracy of segmentation masks. These masks are used for defining regions of interest within an image. Many possible regions of interest exist within an adult *Drosophila* wing blade including the interveins, the longitudinal veins, and the marginal hairs. Semi-automated wing analysis platforms, such as FIJIWings, make use of the trainable Weka segmentation module to identify these regions (Arganda-Carreras et al. 2017; Dobens and Dobens 2013). However, training these programs to identify the regions of interest in the *Drosophila* wing is time consuming and may need to be repeated when using images from different cameras. Alternatively, packages such as WINGMACHINE rely on traditional image segmentation techniques, such as image thresholding, where parameters need to be recalibrated for separate datasets. Additionally, conventional image processing algorithms face a challenge in accurate processing of wing images that might be obtained from variable imaging conditions, such as changes in background lighting or wing rotation. For example, the WINGMACHINE pipeline requires a specific wing orientation for extraction of landmark positions. Recent work assessing wing phenotypes used an open source ML-based pixel classifier ILASTIK (Sommer et al. 2011) for the task of segmenting the overall wing blade (Alba et al. 2020). To date, there is not a fully automated and high throughput image analysis pipeline that can be used for processing a broad range of phenotypes.

Convolutional neural networks (CNNs) serve as an alternate segmentation algorithm that can be adapted to new identification problems (K. G. Kim 2016). As a first step, we retrained the last few neural network layers of a pre-trained U-Net model (Falk et al. 2019; Ronneberger, Fischer, and Brox 2015; Zhou et al. 2018), which is a DL-based image segmentation pipeline for identifying different regions of interest within the adult wing. U-Net was chosen for this task due to the fact that the network relies on data augmentation for efficient use of annotated samples. This process leads to a reduction in the size of the training dataset. In this effort, we used a batch size of approximately 1,000 *Drosophila* wings as the initial training dataset (Kumar et al., in preparation). Details about the images used for training, along with the training implementation, are provided in the methods and SI section S1.

This training process annotates four different regions within an image. These regional classes are: the non-wing background, the intervein regions, the veins, and the periphery hairs (Figure 1B). Training U-Net through PyTorch using a GPU (Ketkar 2017) resulted in a deployable model with an overall accuracy of 95% (Figure 1C’). The U-Net model was trained to be compatible for images either taken using a medical slide scanner or EVOS microscope for RGB channel images at magnification of 4x or higher (Figure 1D). For images at lower resolution and grayscale images, we used the open source ML-based pixel classifier ILASTIK to generate segmentation masks (Sommer et al. 2011). The ILASTIK toolkit extracted 37 features for each color channel within each pixel. These features included intensity, edge-detection, and texture features. Following the extraction step, a random forest classifier from sci-kit learn was used to obtain a consensus classification for each pixel (Pedregosa et al., n.d.). When training MAPPER, ground truth images are added iteratively to reduce the calculated uncertainty of each pixel until the calculated uncertainty reaches a minimum threshold desired by the user (Figure 1E). This segmentation mask is then imported into the custom pipeline of MAPPER for high throughput morphometric quantification of features. Full details on training the image segmentation pipeline are provided in SI section S1.

### MAPPER provides multiplexed analysis tools for high dimensional morphological features

A key feature of MAPPER is the classification of individual intervein region based on geometric and shape features. This is carried out by training a ML-based intervein classifier that takes unlabelled intervein regions from the segmentation mask as an input and classifies them according to their location in the adult *Drosophila* wing (Figure S1). Using these labelled interveins, MAPPER then identifies individual veins, intervein regions, and extracts wing shape features (SI section S3). The systematic labeling of interveins also allows construction of quantified phenomes that can establish geometric similarities and dissimilarities between disparate wing samples.

Segmentation masks generated either by U-Net or ILASTIK are imported into a custom MATLAB pipeline that fills holes, smoothens the edges, and identifies continuous intervein regions. For training a ML-based intervein classifier, morphological features were first extracted for each manually labelled intervein (defined in Figure 2A). EFD-based shape descriptors were first extracted for each intervein to train the classifier (Kuhl and Giardina 1982). The key advantage of using such a framework is that EFDs produce a robust translational and rotation invariant representation of the intervein shape (Kuhl and Giardina 1982).

**Figure 2.**
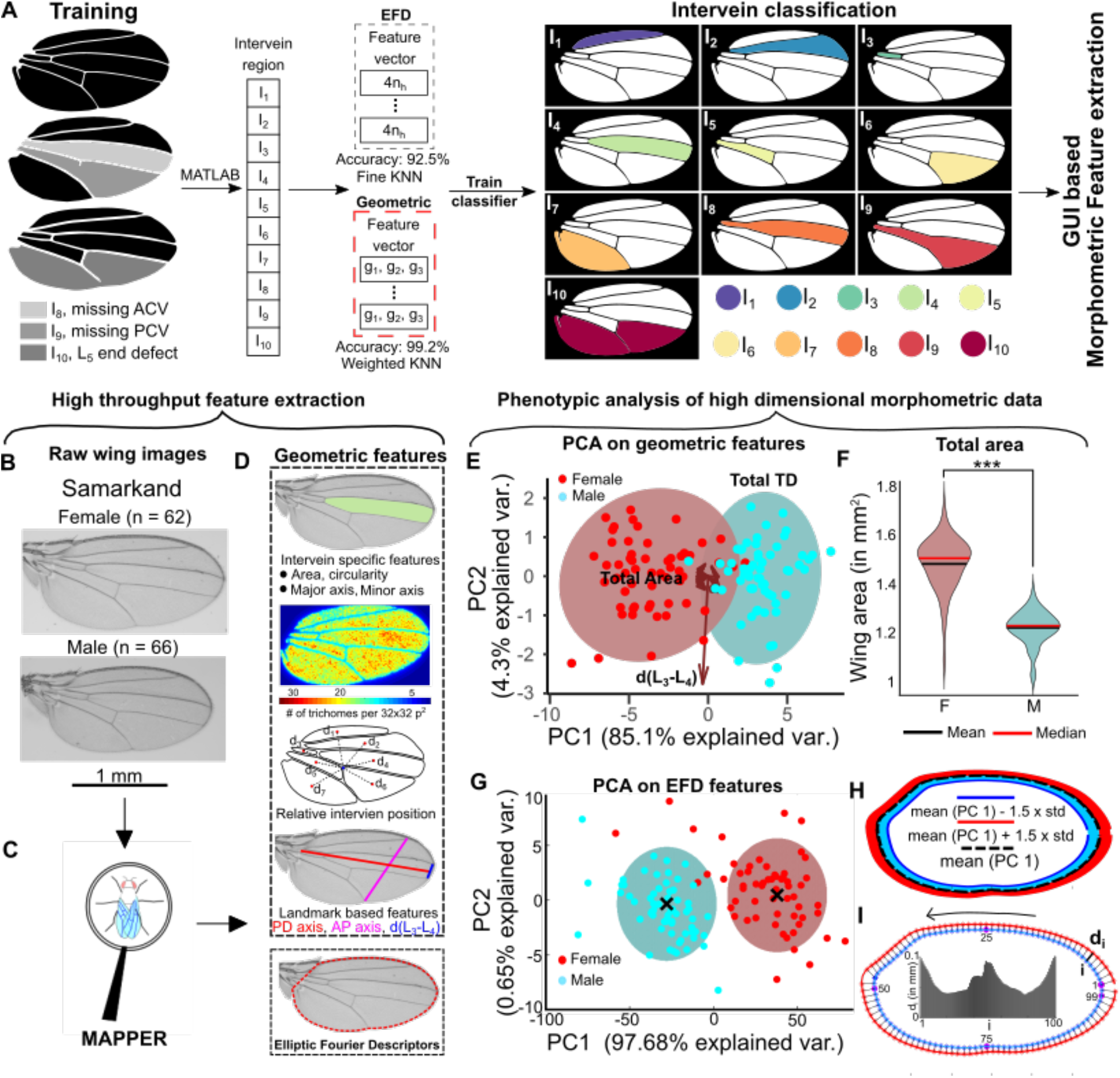
MAPPER automates classification and extraction of an ultrahigh (>100) dimensional morphological feature set. **(A)** During training, each intervein region identified through U-Net was manually labelled. EFD features along with the geometric features were extracted from the labelled intervein regions to train a machine learning (ML)-based classifier. This trained model then identifies the intervein regions based on the input binary mask and the associated features of each intervein region. Edge cases of ACV/PCV and marginal L_5_ defects were included in the analysis. **(B)** Male and Female wings from the *Drosophila melanogaster* Samarkand strain were analyzed to show the phenotypic profiling using an ultrahigh dimensional morphological feature set ((Sonnenschein et al. 2015)). The sample size of each population has been listed on top of wing images. **(C)** MAPPER processed the raw wings. **(D)** MAPPER generates a wide number of geometric features and extracts EFD coefficients fit to the wing margin. **(E)** PCA carried out on the geometric features extracted using MAPPER reveals the largest variance in the data observed in terms of overall wing area and trichome density. Clustering carried out on the first two principal components also revealed two distinct clusters isolated based on features extracted using MAPPER. **(F)** Violin plot showing the distribution of area of male and female wings. Solid red line indicates the median and solid black line indicates the mean of each population. **(G)** PCA on EFD revealed most of the variance in the data concentrated only in the first principal component. A clustering operation also revealed the presence of two distinct clusters representing populations of male and female. **(H)** Standard deviation in the direction of PC1 was calculated for the entire population of wings. PC1 was varied by adding and subtracting 1.5 times the standard deviation along PC1. Reverse PCA was then used to obtain the desired EFD coefficients in which the contours were reconstructed. **(I)** EFD was used to construct mean wing shapes representing the male and female populations. 100 points were sampled from the male wing and their minimum distance from the female wing was calculated to quantify local size differences within the two populations. The variation of size is drawn as a bar graph where the x axis is representative of the points sampled in male wing. Location of points sampled is indicated in the plot.

EFDs are determined by fitting a Fourier series to the periodic function obtained from the closed *Drosophila* intervein contour (Figure S3). The accuracy of an EFD fit varies with the number of harmonics used in the expansion. We fitted EFDs to the seven intervein regions of a wildtype wing to estimate the appropriate number of coefficients required for an accurate representation of shape. The error between the actual contour and the shape approximated by the EFD decreased as the number of terms in the EFD increased (Figure S3B). The first ten terms of the EFD were selected for representing the shape of each intervein.

In addition to the EFDs, we also extracted basic geometrical properties of the intervein regions including: the ratio of an individual intervein area with respect to the area of the whole wing, the circularity of the region of interest (ROI), and the aspect ratio of each intervein region. Extracted EFD coefficients alongside the geometric features of individual interveins were used to train separate models for the task of intervein classification (Figure 2A). This prevents overfitting and selects the set of features that can be used best to classify the interveins. We found that a KNN-based classifier offers the best cross-validation accuracy of the eleven different support vector machine (SVM) and KNN classification methods tested (Figure S4C). The confusion matrix, which defines the accuracy of the classifier, is shown in Figure S4D. Overall, the KNN classifier reported an accuracy of about 92.5% when trained on EFD-based features and 99.2% when trained on the geometric features of each intervein.

Based on this, we used intervein classification scheme based on geometric features for classification of interveins to analyze images (Figure 2A). In summary, for any new segmentation mask, the geometric features described above are extracted from each intervein. The features are then passed into the trained intervein classification model that classifies and labels each intervein region.

The unique labelling of interveins is followed by a series of operations to extract morphological features from the wing blade (SI section S3). The approach also systematically extracts more localized geometric features that can be used for phenomic analysis. One of the key features extracted using MAPPER are the EFD coefficients for the wing periphery (SI section S3D, Figure S6). For this particular step, the EFDs were not normalized against size for quantification of changes in area of the wing. To do so, we modified the original algorithm so that the EFDs produced are sensitive to size changes. This is accomplished by removing the normalization step in previous MATLAB code (Thomas 2020) where the EFD coefficients are normalized by the semi-major axis of the first ellipse (SI section S3). The coefficients of the Fourier series act as additional features, each of them carrying a local shape property. These coefficients can not only be used for screening local shape changes within the wing, but also can be used to estimate an average shape for a particular genotype. Another unique advantage of MAPPER is that it quantifies anatomical axes such as the AP and the PD axis of the wing blade (Figure 2C-D). The labelled interveins are used to delineate the L_2_, L_3_, L_4_ and L_5_ veins along with the cross veins (Figure S5). Identification of veins is also accompanied by quantification of landmark positions within the wing and consequently the anatomical axes. In summary, MAPPER extracts a high dimensional fingerprint consisting of *Drosophila* wing specific morphological features in a high throughput manner.

### Representative statistical approaches for phenotypic analysis

In this section, we describe a series of statistical techniques for probing and analyzing the variation in phenotypic traits within and between wing populations of wings. For this analysis, we processed 128 adult wing images of *Drosophila melanogaster* from the Samarkand strain using the GUI-based interface of MAPPER (Figure 3, Figure S10) (Sonnenschein et al. 2015).

**Figure 3.**
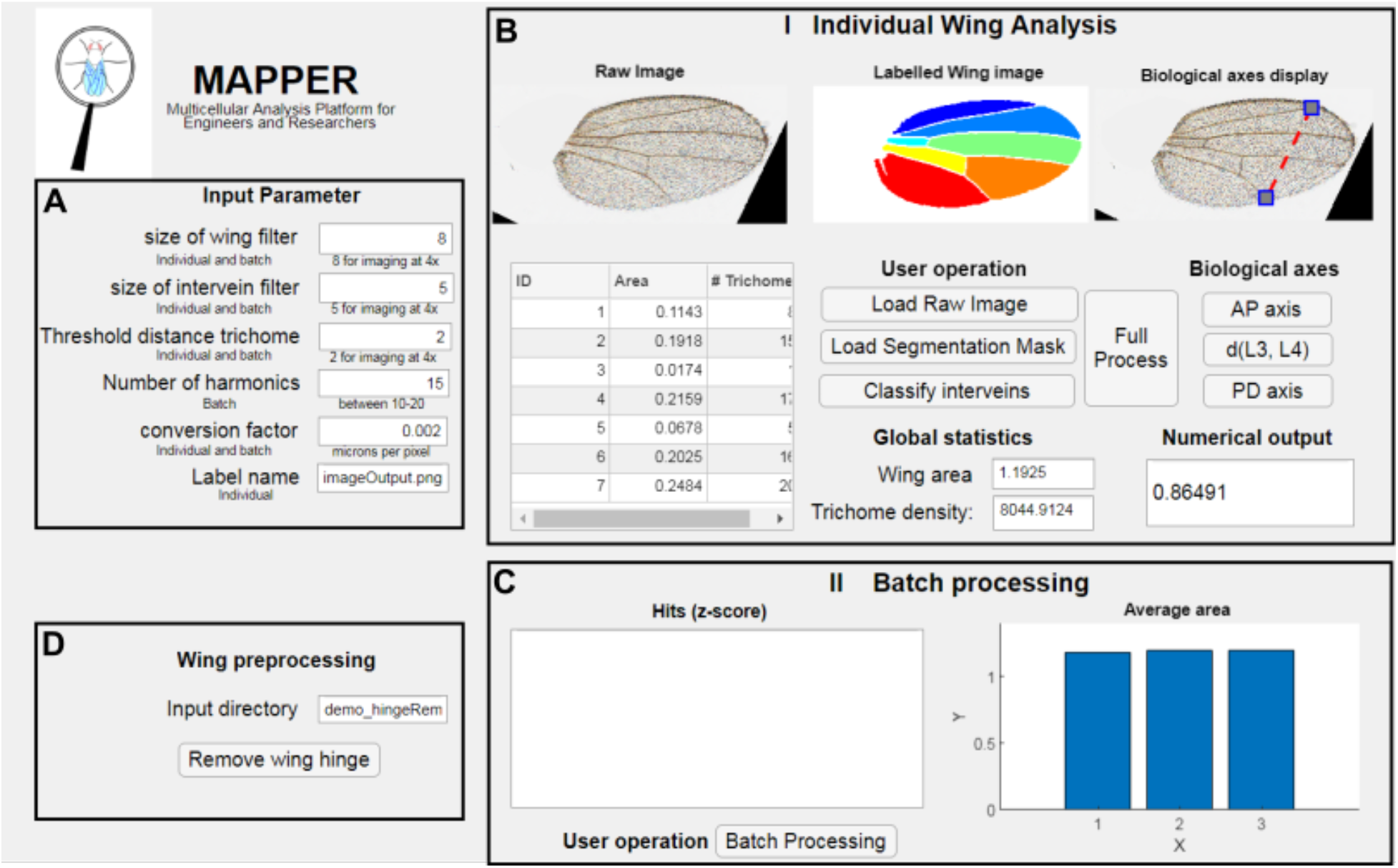
MAPPER application. he MATLAB MAPPER application allows for individual. **(A-B)** and batch **(D-C)** processing of adult *Drosophila* wings. Details about the app interface and its use can be found in the SI text.

The high dimensional morphometric fingerprint generated by MAPPER separates the male and female population of wings (Figure 2E,G). The sex-based difference in patterning and size of adult *Drosophila melanogaster* wing is known and well studied (Testa and Dworkin 2016). We also show that the EFD-based features can be used to probe into this sex-related shape changes more locally (Figure 2H). To highlight this utility of EFDs, it was separately analyzed from the geometric features. First, Principal Component Analysis (PCA) (Wold, Esbensen, and Geladi 1987) carried out on the geometric features revealed that the maximum variance within data was distributed majorly between the first two principal components (89.4%) (Figure 2E, Figure S10C-D). Analysis of the loadings for the first principal component showed that overall wing blade area explains the majority of major variance within the data.

As expected, the area of the female wing was significantly larger than a male wing (*p*-value < 0.001) (Figure 2F). A plot of the first two principal components shows the two distinct clusters of male and female populations (Figure 2E). Further, PCA applied on the EFD coefficients revealed a total variance of about 97% distributed along PC1 alone (Figure 2G). The observed variance along PC1 is attributed to the known overall size differences between the male and female population of wings (Figure 2H). Clustering carried out on the first two principal components using Gaussian Mixture Models (Yang, Lai, and Lin 2012) (GMM) was also able to distinguish the male and female population of wings (Figure 2G). The mean shapes of each cluster can also be used to highlight shape changes between the male and female populations more locally. We used the mean contours of each population and measured variation peripheral growth along the normal direction (Figure 2I). There was more growth along the PD axis as compared to the AP axis, which is necessary for maintaining a uniform scaling of these anatomical axes with overall size of wing blade (Figure S11). This uniform scaling also confirms that the normalized length of the AP na DV axis is equal for both the male and female wings. We explored these scaling relationships in more detail in the first case study conducted on wings belonging from four species of *Drosophila*. Details about the statistical techniques followed in analysis of this dataset are included in the SI (SI section S5).

In summary, the high dimensional feature set generated by MAPPER distinguishes different populations of wing morphologies or shapes. Implementing these approaches demonstrates size and shape differences between male and female populations of *Drosophila* via systematic analysis of features extracted. In particular, females had significantly larger wings (*p*-value < 0.001) consistent with sexual dimorphism of insulin signaling activity (Testa and Dworkin 2016) while males had significantly larger trichome densities (*p*-value < 0.001). We also show how variable regulation of growth is necessary for maintaining perfect scaling of anatomical axes such as the AP axis and PD axis of the wing. This analysis was done despite the wing images being imaged at different rotations via the MATLAB-based MAPPER application.

### MAPPER provides fast, precise segmentation and extraction of wing morphologies

We validated MAPPER’s performance through a systematic comparison of metrics such as wing blade area and trichome density. To do so, we benchmarked MAPPER and FIJIWings with a data set of wings of varying size generated by genetically perturbing insulin signaling. Insulin and insulin-like growth factors regulate metabolic activity (Belfiore et al. 2009; Kurtzhals et al. 2000; Samani et al. 2007). Dysregulation of insulin signaling causes a variety of human diseases including diabetes, insulinoma, metabolic syndrome, ovary syndrome, and auto-immune disorders (Dunaif et al. 1989; Kahn, Hull, and Utzschneider 2006; Wang et al. 2017). In *Drosophila,* the InsR homolog regulates cellular proliferation (Brogiolo et al. 2001). Loss of function of InsR in wing imaginal discs reduces final wing size (Chen, Jack, and Garofalo 1996).

As expected, quantification of wing size and trichome number shows that activation of InsR signaling increases wing size and trichome number. Conversely, suppression of InsR signaling reduces wing size and trichome number (Brogiolo et al. 2001). The overall area of wings along with their trichome densities were comparable when they were measured through FIJIWings and MAPPER (Figure 4B, Figure 4C). FIJIWings tended to over-segment tissues as regions containing marginal wing hairs were often misclassified as intervein regions (Figure 4A). This was not observed in any of the segmentation masks produced by MAPPER. Additionally, MAPPER processed images faster than FIJIWings by two orders of magnitude, thereby enabling high throughput screening applications (Figure 4D). The areas estimated by each tool, respectively, were comparable with manual measurements. A line with slope of 1.03 ± 0.0041 (R^2^: 0.99) was obtained for the fit of MAPPER measurements versus measurements by hand, confirming the accuracy of MAPPER (Figure 4E).

**Figure 4.**
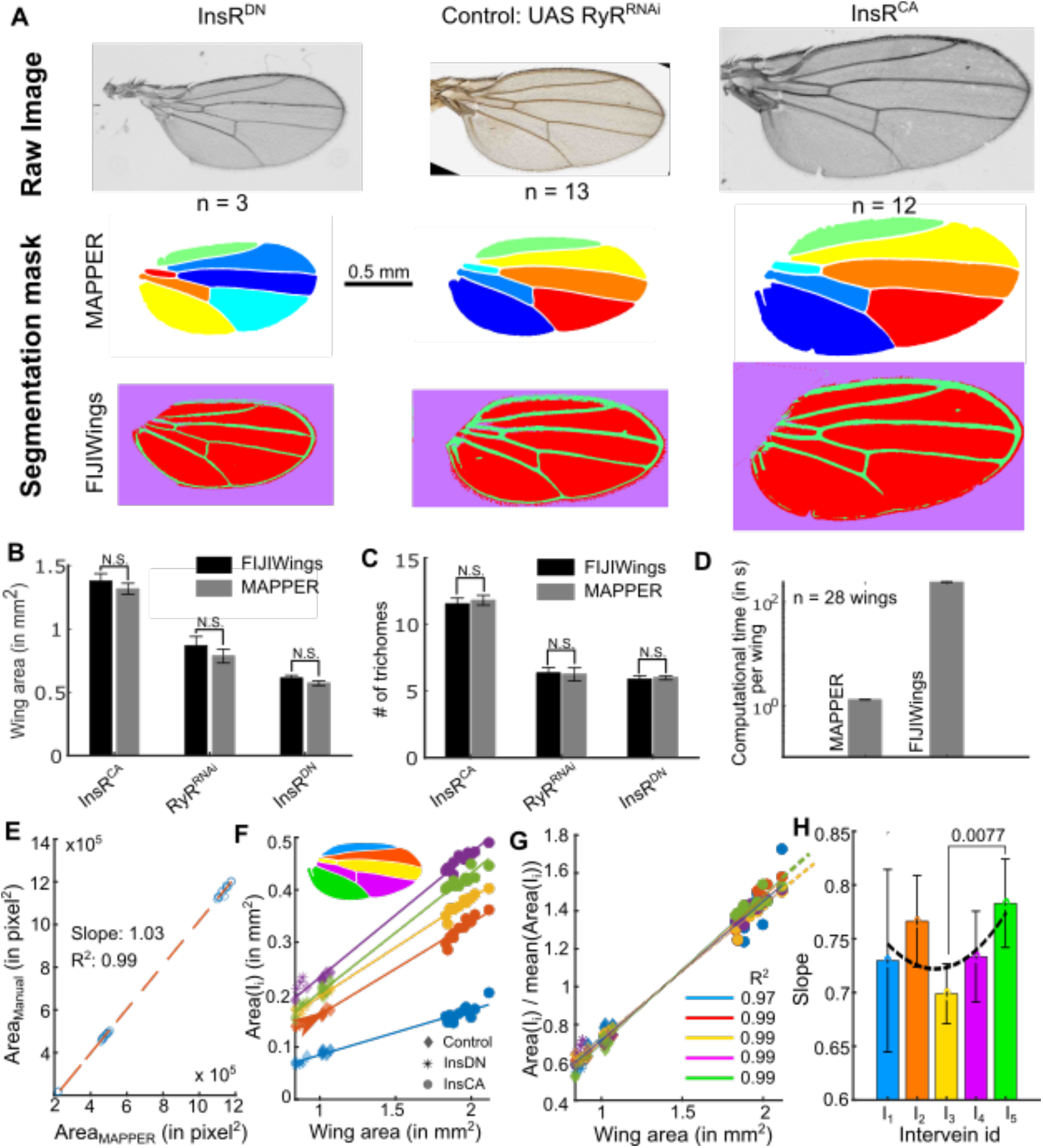
MAPPER provides precise segmentation and extraction of wing morphology. **(A)** MAPPER automates identification and labeling of the individual intervein components with high accuracy. InsRDN is the dominant negative form of insulin receptor and InsRCA is the constitutively active insulin receptor. RyRRNAi is the GAL4-UAS knockdown of the ryanodine receptor, which is not expressed in the wing disc. Full genotypes are the MS1096-Gal4 lines crossed to each of the indicated UAS lines. **(B)**, **(C)** Comparison of wing area and number of trichomes, respectively, as calculated by MAPPER and FIJIWings. Error Bars indicate standard deviation in measurements. **(D)** The computational time of MAPPER is two orders of magnitude faster than FIJIWings. **(E)** Measurements of wing blade area done by MAPPER versus measurement of wing blade area done by hand. The slope and R2 values indicate that MAPPER produces identical values to those produced by hand. **(F)** Perturbations in InsR lead to variation of area of intervein regions when scaled to overall change in wing blade area. Linear regression was used to fit straight lines that show the scaling relationships for individual compartments. Line color corresponds to the intervein regions in the inset of the graph. **(G)** Variation of area of each intervein region scaled by mean area of the population with overall change in area of the wing blade. **(H)** Slopes of lines fit to data points in G. This clarifies the role of insulin signaling in maintaining proper scaling relationships between intervein regions. N.S. indicates no statistical difference between comparisons.

Further, MAPPER distinguished and classified the individual intervein regions to output geometrical statistics for the labelled components. This feature extraction capability is unique to MAPPER. This new capability allowed us to explore how individual intervein regions scaled with an overall increase or decrease in wing area resulting from perturbations in the insulin signaling pathway. In particular, we observed differences in scaling factors corresponding to changes in intervein area I3 and I5 (*p*-value = 0.0077) with respect to change in overall wing blade area (Figure 4H). This result indicates a possibility of differential regulation of the insulin signaling pathway in controlling growth within individual regions of *Drosophila* wing imaginal discs. This analysis was carried out by first plotting the area of each intervein region against the overall wing blade region. A straight line was then fit to these data points (R^2^ of fits included in Figure 4G). High values of R^2^ indicate that the growth of each intervein was linear in response to overall growth. To evaluate the percentage growth or size change of individual interveins, we also re-evaluated the scaling relationships by normalizing the area of each intervein with the overall area of wing blade (Figure 4G). Comparing the slope of the fit lines reveals a minima in the degree of scaling, for the normalized data, as one moves towards the center of the wing from I1 and I5 towards I3 (Figure 4H). This might indicate that intervein area scaling depends on the morphogen regulated transcription-factors defining intervein regions in the wing. The existence of this gradient in regional scaling suggests coupling of insulin signaling regulatory activity with tissue patterning function. Taken together, these results not only validate the accuracy and speed of MAPPER, but also highlight how its improved methodology can discover high-dimensional scaling relationships under a broad range of genotypic perturbations.

### *Case Study* I: MAPPER reveals species-specific differences in wing size and pattern

We used MAPPER to quantify morphometric phenotypes of wings in four species: *D. melanogaster*, *D. simulans* (a species in the *melanogaster* subgroup), *D.ananassae* (a species in the *melanogaster* group), and *D. virilis* (a species outside the *melanogaster* group (Lage et al. 2007). We measured the total wing area in male and female individuals in each of these species (Figure 5A). Wing area was revealed to be larger in females when compared to males in *D. melanogaster, D. simulans*, and *D. ananassae*. However, in *D. virilis* adult wings are larger in males than in females (Figure 5A,B). Next, we examined vein patterns. While wings in all species have the same venation pattern (Figure 5A), as revealed by statistically similar intervein areas relative to total wing area (Figure S8A), we found size-independent differences among species, especially in females (Figure S8A). More specifically, by measuring two different segments of the longitudinal vein L_5_ (Figure 5C), we observed that the relative location of the posterior cross-vein that connects the longitudinal veins L_4_ and L_5_ is approximately the same for *D. melanogaster* and *D. simulans*, but is located more distally in *D. ananassae* and *D. virilis* (Figure 5C’, Figure S8B). We also found species-specific differences in the relative areas of anterior (A) and posterior (P) regions of the wing (Figure 5D). In *D. simulans* and *D. ananassae* the A and P regions are approximately of the same size, but in *D. melanogaster* and *D. virilis* the P region is about 10% and 20% larger than the A region, respectively (*p*-value < 0.001) (Figure 5D’, Figure S8C). Details about the tests for determining the statistical significance of these comparisons can be found in the SI section S6 while individual *p*-values are listed in Figure S9. It should be noted that the dataset analyzed for this analysis did not have sufficient resolution for analysis of trichome density patterns.

**Figure 5.**
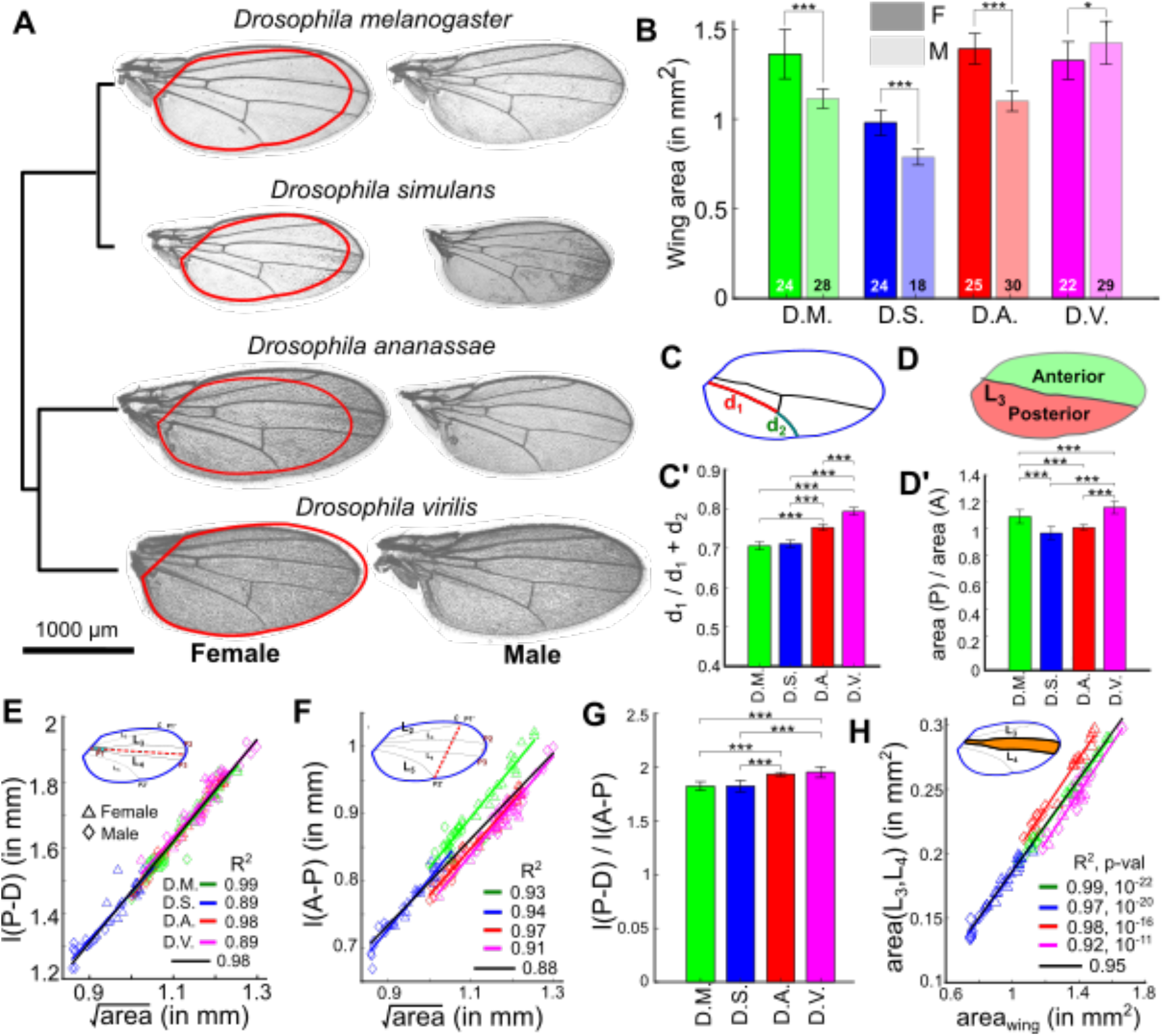
MAPPER identifies differences in wing size and patterns among *Drosophila* species. **(A)** Representative male (right) and female (left) wings of four different species. Red contours on female wings represent the outline of the corresponding male wing. The dendrogram in the left is representative of hierarchical clustering based on different features extracted using the pipeline. **(B)** Quantification of wing areas by MAPPER for wings from different species and different sexes. **(C-C’)** Quantification of shift in posterior cross vein position in female wings (d_1_ is defined as the segment of L_5_ from the proximal end of the vein to posterior cross vein, d_2_ is defined as the segment of L_5_ from the posterior cross vein to the distal end of L_5_). **(D-D’)** Relative anterior (A) and posterior (P) areas in female wings. **(E)** Scaling relationships between the length of the proximal-distal (l(P-D)) axis and the overall wing blade area for the various species. Legends for different sexes have been included. Straight lines were fit to estimate the existence of scaling relationships for the four species. R2 of the fit line are reported in the inset. **(F)** Scaling relationships between the length of the anterior-posterior (l(A-P)) axis and the overall wing blade area for the four species. Straight lines were fit to estimate the existence of scaling relationships for the different species. R2 values of the fit line are reported in the inset. **(G)** The l(P-D) to l(A-P) ratio for females from different species. **(H)** Scaling relationships between the area of the intervein region between veins L_3_-L_4_ and the overall wing blade area for the four species. Straight lines were fit to estimate the existence of scaling relationships for the different species. R2 values of the fit line are reported in the inset. (Refer Figure S9 for *p*-values corresponding to all comparisons. Details on the statistical tests done can be found in the section S5 of the SI. * *p* < 0.05, ** *p* < 0.01, *** *p* < 0.001.)

### *Drosophila* wings show a conserved and robust interspecies scaling relationship along their PD axis, but not along the AP axis

We further examined how the PD and AP axes scale within and across species. Along the PD axis, all species follow a similar linear scaling relationship with respect to the square root of the total wing area (Figure 5E), suggesting that there is strong selection in maintaining a proportional PD length across species. Along the AP axis, we also observe a linear scaling relationship for the AP length, but the slopes vary from species to species (Figure 5F). To further explore these results, we plotted the ratio of the PD and AP axes lengths and found that *D. ananassae* and *D. virilis* have a slight but significantly larger ratio than *D. simulans* and *D. melanogaster* (Figure 5G, Figure S8D). Taken together, these data suggest that the length of the AP axis in *D. ananassae* and *D. virilis* is significantly shorter compared to that of *D. simulans* and *D. melanogaster*. Since we see changes between these pairs of species, but not within the species of each pair, our results suggest that the length of the AP axis evolved with the phylogenetic split between these pairs of species (Figure 5A).

In *Drosophila*, the AP axis is patterned by two morphogens: Hedgehog (Hh) and Decapentaplegic (Figure 1A) (Dpp) (Blair 2007). Hh patterns the most central region (L_3_-L_4_ veins) (Vervoort et al. 1999)(Mohler et al. 2000)), whereas Dpp patterns the positions of L_2_ and L_5_ (Affolter and Basler 2007; Restrepo, Zartman, and Basler 2014). To pinpoint whether the changes we see along the AP axis could be attributed to any of these signaling pathways, we compared the L_3_-L_4_ intervein area in these species (Figure 5H). We found that these similarly scale in all four species, suggesting that it is unlikely that these differences are due to variations in the regulation of the Hh signaling pathway. Interestingly however, the areas comprising veins L_2_ and L_5_ with respect to the wing margin appear to scale differently across species (Figure S8A), suggesting that Dpp signaling dynamics diverged during speciation to modulate the proportions of these wings along the AP axis. As a prediction for future studies, these results are suggestive that Dpp transport and/or transduction is variable while Hh is not across species. In sum, MAPPER proved to be a powerful toolkit for identifying new morphogenetic relationships across *Drosophila* species.

### *Case Study* II: The mechanosensitive Ca2+ channel *Dmpiezo* regulates left-right size symmetry in adult *Drosophila* wings

In an ongoing parallel study, we have been screening genetic candidates that may be novel modulators of wing morphogenesis (Kumar et al., in preparation). Here, we highlight the use of MAPPER in identifying surprising and interesting insights into the roles of Ca^2+^ channels in regulating wing sizes. Previous studies found a single homologue to the mammalian *Piezo1* in *Drosophila* (Coste et al. 2012). We used the GAL4-UAS binary gene expression system to either knockdown or overexpress *Dmpiezo* channels via a wing imaginal disc tissue-specific GAL4 driver. We also characterized the *Dmpiezo* knockout line (BDRC #58770) for a comparison with the GAL4-UAS mediated perturbations. Interestingly, *Dmpiezo* wings did not show any severe phenotypic traits (Figure 6A). Additionally, the mean area of wings resulting from *Dmpiezo* knockout remained unchanged when compared with the parental lines used for generating the knockout. Surprisingly, the *Dmpiezo* KO wings showed a significant increase in the within population variance of wing sizes (*p*-value < 0.05) and dysregulated left-right symmetry in size of the sister wings (Figure 6F,G,I). The difference in the sizes of sister wings was normalized by the mean wing area of the whole population. A knockout of *Dmpiezo* resulted in a higher incidence of cases where the difference between the areas of sister wings was significantly larger than the control parental lines, with multiple instances of large asymmetries between sister wings (Figure 6I).

**Figure 6.**
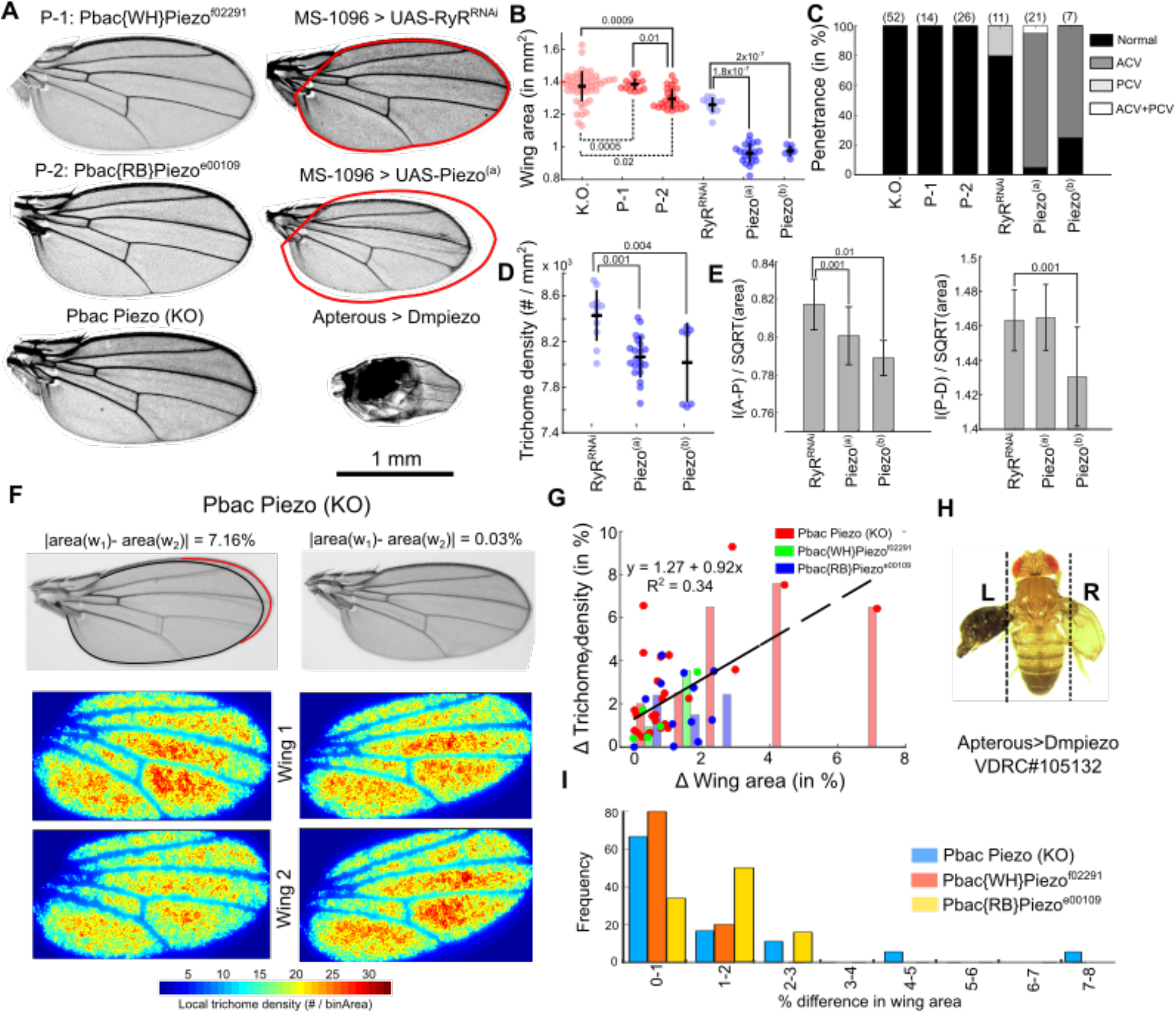
Piezo regulates left-right symmetry along with growth and patterning of wings. **(A)** *Drosophila* wings from multiple genetic perturbations related to *Piezo*. **(B)** Beeswarm plot showing distribution of wing area for different perturbations to Piezo. KO represents the *Dmpiezo* knockout line while P-1 and P-2 are the parents used to generate the knockout line. Piezo^(a)^ and Piezo^(b)^ are the two commercially available overexpression lines for Piezo. The overexpression was carried out using a MS-1096 wing specific Gal4 driver. UAS-RyR^RNAi^ crossed with the MS-1096 Gal4 driver serves as the control for these comparisons. p-value over solid lines indicate statistical significance for comparison of means while the p-values written over dashed lines are for comparing variances of these populations. **(C)** Quantification of phenotypic penetrance of vein defects. Penetrance is the measure of percent of a particular phenotype in a given group. (ACV: Anterior cross vein, PCV: Posterior cross vein) **(D)** Beeswarm plot showing distribution of trichome densities for different perturbations to Piezo. **(E)** Quantification of scaled Anterior-Posterior and Proximal-Distal axes for the Piezo overexpression lines. **(F)** Heat maps showing trichome density patterns of the sister wings for Piezo knockout lines. These heatmaps show that wings with greater difference in area show a correspondingly higher difference in trichome density. **(G)** Plot showing correlation between difference in area and trichome densities. **(H)** Adult ap>Piezo^RNAi^ fly showing significant patterning and symmetry defects. **(I)** Bar graph showing distribution of differences of area between sister wings of the Piezo knockout versus knockdown lines.

Interestingly, MAPPER also revealed that these differences in the area of sister wings correlates with the differences in the trichome densities (Figure 6F, Figure 6G). Taken together, these results suggest that *Dmpiezo* has a significant role in regulating the left-right asymmetry and robustness in organ size growth of *Drosophila* wings. To further test whether Dm*piezo* regulates bidirectional symmetry in patterning of adult *Drosophila* wings, we knocked down the mRNA expression of *Dmpiezo* in the dorsal compartment of the wing imaginal disc with Apterous GAL4>*Dmpiezo RNAi* (BDRC#105132). RNAi inhibition of *Dmpiezo* led to an adult with severe, but different morphological defects in sister wings (Figure 6H). Surprisingly, the severity of phenotypic traits was different between the knockout versus the knockdown lines. This suggests a possibility of genetic compensation where a similarly functioning gene takes up the role of the downregulated component (Song et al. 2019). Another explanation for these differences is a possible RNAi mediated off target effect. However, BDRC does not report any predicted off target effects with the UAS-*Dmpiezo* RNAi line used. We also used two commercially available overexpression lines (BDRC #58772, BDRC#58773) to overexpress these receptors in the wing imaginal disc pouch. An overexpression of *Dmpiezo* channels led to a reduction in overall wing blade area. Both the overexpression lines resulted in adult wings with nearly 25% reduced size (p-value < 0.001) as compared to the MS1096>RyR^RNAi^ control (Figure 6A). The reduction in wing size was also accompanied by high incidences of missing anterior cross veins (Figure 6A, C).

Currently, the exact mechanism by which Piezo-stimulated calcium signaling regulates robustness in growth and patterning is still unclear. However, these results indicate a novel role of *Dmpiezo* in regulating *Drosophila* wing development and consequently epithelial morphogenesis. Here, MAPPER provided a high throughput and accurate determination of the size and trichome densities of hundreds of wing images in a very short amount of time. This type of analysis would not have been feasible without MAPPER’s automated batch processing capabilities and fast computational speeds.

## Discussion

### Features and strengths of MAPPER

Conventional image processing techniques are often unable to process images of model organism morphologies that have been generated with different imaging systems. For example, traditional image processing pipelines have difficulty analyzing images taken with multiple different lens objectives, lighting conditions, or rotational orientations. As such, these pipelines often fail at accurately processing images beyond the initial dataset for which it has been developed.

Additionally, the number of morphometric features that are extracted are low dimensional, making them unsuitable for detecting subtle quantitative changes. MAPPER supersedes previous pipelines by using a statistical learning framework with the latest computer vision and machine learning approaches. A key feature of MAPPER is its hybrid, modular framework. The first component is a DL-based pixel classification module that segments individual regions of wings. The second module labels each intervein region according to its shape-based features. In conjunction, these individual pipelines allow MAPPER to generate a wide variety of geometrical and pattern-based features of wing images. Further, using the statistical learning algorithms in the pipeline also help in increasing the overall computational speed in processing wing images. The precise labelling of interveins allows for reconstruction of veins and automated extraction of landmark-based measurements, such as the AP and PD axes. In summary, the coupling of two modules with an integrated diverse class of functions, automates the systematic generation of high dimensional geometric and pattern-based features for a large volume of wing image data.

### Implications of insights and new hypotheses generated by MAPPER

Our analysis of adult wings in different *Drosophila* species using MAPPER reveals two key observations. First, we noticed a reversal in sexual dimorphism when comparing species within the *melanogaster* group with *D. virilis*. Particularly, wings of *D. melanogaster*, *D. simulans*, and *D. ananassae* are larger in females than in males. However, in *D. virilis* the opposite phenotype is observed (Figure 5A-B). How wing size is differentially regulated in a sex-specific manner across species is unclear, but our data suggest that the dimorphism that makes female wings larger than male wings arose at some point in the divergence between the *melanogaster* and *virilis* groups. Second, the length of the wing PD axis across species and sexes shows a conserved scaling relationship with respect to wing size (Figure 5E), suggesting that while ecological and genetic changes may exert pressure on overall wing size, preserving a scaling relationship between length of the PD axis and total wing area in all species may be essential. In contrast, the AP axis in *D. ananassae* and *D. virilis* is smaller with respect to what would be predicted from the scaling relationship of *D. melanogaster* and *D. simulans* (Figure 5F). Since BMP/Dpp signaling is responsible for patterning and growth along this axis, we predict the variation in this pathway between species can explain the larger AP axis in *D. melanogaster* and *D. simulans.* Variation in Dpp pathway activities between species may also explain why the posterior cross vein is located more distally in *D. ananassae* and *D. virilis* compared to *D. melanogaster* and *D. simulans* (Figure 5C-C’). This is because the specification of the posterior cross vein depends on pupal BMP signaling driven by Dpp and Glass-bottom-boat (Gbb) ligands (Ray and Wharton 2001).

Bidirectional symmetry in growth and patterning of the left and right axis of development has received significant attention in vertebrates and higher level of organisms (Martindale and Henry 1998; Holló 2015), but the pathways and mechanisms through which this symmetry is maintained during morphogenesis is still unclear. Here, the high throughput processing capabilities of MAPPER led to discovery of a surprising new role of *Dmpiezo* in regulating symmetrical and robust wing size control. Previous studies have established Piezo, a mechanosensitive Ca^2+^ channel, as one of the key regulators of epithelial morphogenesis in mammals (Eisenhoffer et al. 2012; Gudipaty et al. 2017). Surprisingly, knockout of *Dmpiezo* in *Drosophila* leads to viable adults with no qualitative morphological phenotypes. Surprisingly, MAPPER enabled us to determine that loss of function of *Dmpiezo* in the adult fly significantly increased variability in wing area compared to the parental lines used to generate the knockout (Figure 6A). The striking variability phenotype is reminiscent of cell competition phenotypes (Brás-Pereira and Moreno 2018). Additionally, *Piezo* regulates cell extrusion and proliferation in a developing zebrafish tail fin (Eisenhoffer et al. 2012; Gudipaty et al. 2017). Based on this, we hypothesize that *Piezo* regulates robustness in cell cycle progression, thereby regulating the robustness in the patterning and the final size of the organ. Adding to this, MAPPER was also used to determine that the difference in final wing blade size between sister wings was greater than 1% for approximately one quarter of all flies with homozygous loss of *Dmpiezo*. One explanation for this observation lies in the fact that *Piezo* induces both proliferation and extrusion (Piddini 2017). Additionally, mutations in the gene Hid, a regulator of cell competition induced cell death, increases bilateral asymmetry between the adult wings of *Drosophila* (Neto-Silva, Wells, and Johnston 2009). Ultimately, the ability for *Piezo* to regulate both cell extrusion and cell proliferation would be biologically beneficial to an organism because malfunction of *Piezo* would not result in a strong directional effect towards either larger or smaller tissue size in most circumstances (Piddini 2017). *Piezo*, therefore, can be inferred to be needed in correcting improper development.

Additional results obtained by MAPPER suggest that wing-specific knockdown of *Dmpiezo*, via mRNA degradation, results in a reduction of final wing size (*p*-value < 0.001, Figure 6B). This poses an interesting question: what is the mechanism that causes the reduction of final wing size with downregulation of *Dmpiezo*? One of the possibilities is that *Piezo* may be responsible for regulating cell proliferation. Loss of *Dmpiezo* can result in a loss of stretch induced mitosis, thus reducing the overall size of wing. Integrated calcium intensity over time is another factor that has been established to be correlated with final organ size (Brodskiy et al. 2019). Hence *Dmpiezo* may also be responsible in regulating final organ size through a regulation of intracellular calcium dynamics. A key question for future investigation is resolving -the difference in severity of phenotypes between the *Dmpiezo* knockout and RNAi induced Dmpiezo knockdown lines. The adult wings in a complete *Piezo* knockout line looked qualitatively normal, while RNAi mediated knockdown of Piezo resulted in wings with severe phenotypic defects. This difference may be attributed to genetic compensation as a result of the total knockout of the *Piezo* gene. Previous studies also suggest that genetic compensation counteracts the effects of a knockout of *Piezo* in neural cells (Song et al. 2019). Many questions about the relationship between *Piezo*, calcium signaling, and regulation of the cell cycle still remain. Currently, the mechanism by which *Piezo* stimulated calcium signaling can result in both cell extrusion, as well as cell proliferation remains unclear.

### Current limitations and future extensions

The new findings from these case studies demonstrate that, in the current age of big data phenomics, manual phenotypic characterization provides an incomplete characterization of phenotypic variation in samples. Here we present a novel, hybrid ML-based approach that was used to automate high throughput processing of adult *Drosophila* wings. The image segmentation capabilities of MAPPER can be easily extended to any insect wing by using training datasets from different imaging sources and multiple insect species. One of the greatest strengths of MAPPER is its automated intervein classification module. In the future, MAPPER can also be adapted to perform phenotypic analysis at a whole organism level. Recently, there have been several attempts to extend depth of field and multiview imaging of insects (Ströbel et al. 2018). The advancements in the field of DL-based smartphone imaging has allowed smartphones to be used for the acquisition of multiview datasets. Integration of algorithms such as Multi-View Deep Extreme Learning Machine (MVD-ELM) can easily be used for the task of 3D segmentation of specific organs (Ahmed et al. 2018; Xie et al. 2015). In summary, MAPPER provides new capabilities to rigorously fit form to functions for a broad range of applications ranging from comparative genomics, drug target discovery, and phenotypic screening.

## Experimental Procedures

### Fly culture, wing collection, and imaging

Wing-specific GAL4 drivers were grown at 25oC. Virgins were collected twice a day from the bottles. Virgins were crossed with males that carry the indicated UAS-TRiP line constructs in a ratio of female:male of 10:3. Adult flies were harvested within 7 days of enclosure and wings were removed. Wings were mounted on microscopy slides to obtain high resolution images (analysis has been extended to all of the wings: 360 wings analyzed in different case studies for this paper). Wings were placed in ethanol, and approximately 15 wings were mounted on each slide in Permount medium (Fisher Scientific, SP15) using standard procedures. For the case study I involving wings from four Drosophila species, the wings were incubated overnight in 70% ethanol.For the case study related to InsR, slides were batch-imaged using an Aperio (Leica) slide scanner at 5X and 20X magnification (Courtesy of South Bend Medical Foundation). Slide scans were obtained in SVS format which allows rapid visualization of large, high-magnification images at various scales. To further analyze the data, these files were converted into TIF format using reaConverter 7 Standard (reasoft). Images wersegmented using ILASTIK. A custom MATLAB script was used to identify the position and orientation of wings. Individual wings were rotated and flipped to the canonical orientation (Top: Anterior, Bottom: Posterior, Left: Proximal, Right: Distal), and then the edges of the image were cropped. The size of the cropping box was determined using the dimensions of the ILASTIK generated segmentation mask. Wings of different Drosophila species (case study I) were imaged using a Nikon Eclipse Ci-S microscope using a Jenoptik ProgRes^®^ monochromatic camera and the ProgRes^®^ Capture Pro 2.9 software. Wings analyzed for case study II were imaged using an EVOS microscope.

### Fly lines

| Fly lines | Source | Stock # | Genotype |
| --- | --- | --- | --- |
| MS1096-GAL4, UAS Dcr2 | BDRC | 25706 | w[1118] P{w[+mW.hs] =GawB}B x[MS1096]; P{w[+mC]=UAS-Dcr-2.D}2 |
| Nub-GAL4, UAS-GCaMP6f | (Brodskiy et al. 2019) | N/A | nub-GAL4,UAS-GCaMP6f/CyO |
| Dmpiezo | VDRC | 105132 | P{KK101815}VIE-260B |
| UAS-piezo::GFP | BDRC | 58772 | w[*];P{w[+mC]=UAS-Piezo.GFP}IIA-2/CyO;MKRS/TM6B, Tb[1] |
| UAS-piezo::GFP | BDRC | 58773 | w[*]; sna[Sco]/CyO; P{w[+mC] = UAS - Piezo .GFP}IIIA/TM6B, Tb[1] , |
| Pbac Piezo | BDRC | 58770 | PBac{w[+mC]=RB5.WH5}Piezo[KO] |
| UAS-Rya-r44F RNAi | BDRC | 31540 | y[1] v[1]; P{y[+t7.7] v[+t1.8]= TRiP.JF01100 }attP2 |
| UAS-Insulin Receptor (dominant negative) | BDRC | 8248 | y[1] w[1118]; P{w[+mC]=UAS-InR.del}2 |
| UAS-Insulin Receptor (constitutively active) | BDRC | 8252 | y[1] w[1118]; P{w[+mC]=UAS-InR.K1409A}2 |

#### Fly lines used for case study I

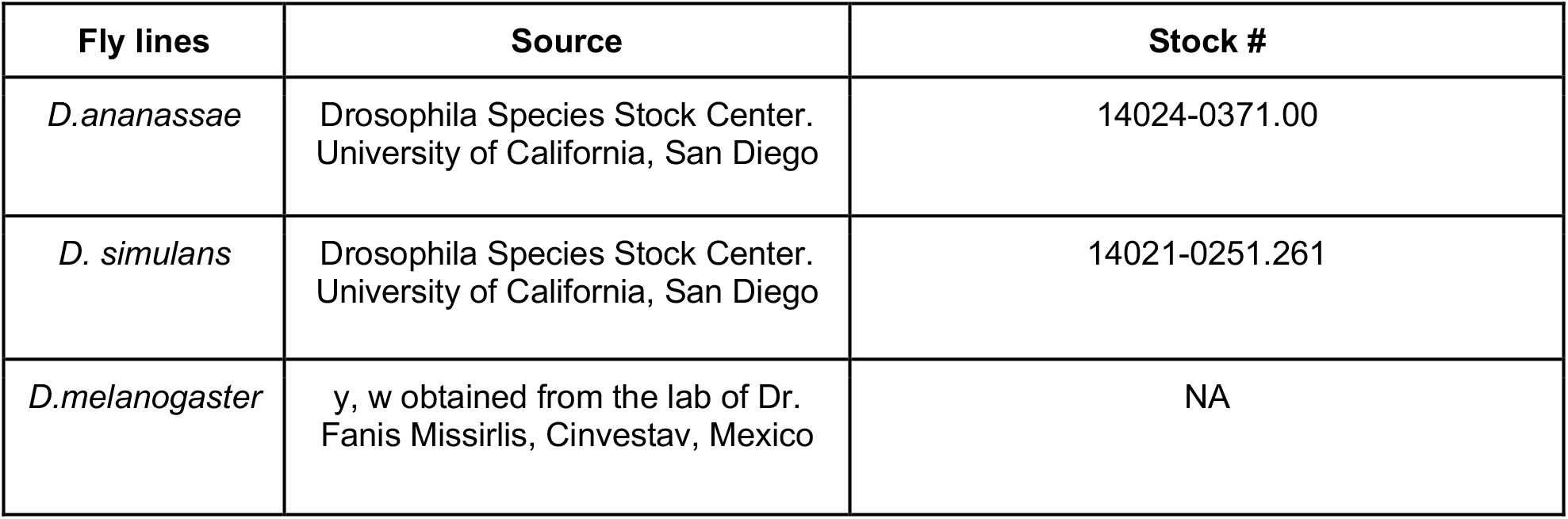
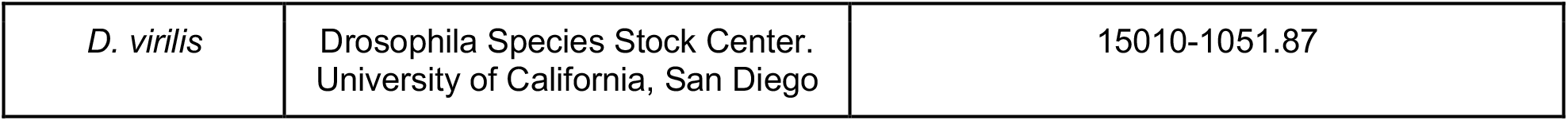

## Acknowledgements

We would like to thank the South Bend Medical Foundation for generous access to their Apero Slide Scanner. We would like to thank Dr. Ramezan Paravitorgabeh, Vijay Kumar Naidu Velagala, Megan Levis, and Dr. Qinfeng Wu for technical assistance and scientific discussions related to the project. The work in this manuscript was supported in part by NIH Grant R35GM124935, NSF award CBET-1553826, NSF-Simons Pilot award through Northwestern University, the Notre Dame International Mexico Faculty Grant Program, and grant CB-014-01-236685 from the Concejo Nacional de Ciencia y Tecnología of Mexico.

## Conflict of Interest

The authors declare no conflict of interest.

## Supplementary Information

### Table of contents

S1 Deep learning-based image segmentation
  S1A U-Net training details: MAPPER utilizes U-Net for training
  S1B Deployment of a trained deep learning model on raw wing image data
S2 Training the intervein classifier
  S2A Generating training data
  S2B Feature extraction for training
  S2C Training a machine learning-based intervein classifier
S3 Morphological feature extraction using the labelled interveins
  S3A Intervein geometric features
  S3B Trichome density of interveins
  S3C Landmark position-based features
    S3C-I Length of proximal-distal axis
    S3C-II Length of anterior-posterior axis
    S3C-III Distance between veins L_3_ and L_4_
  S3D Elliptic Fourier Descriptors (EFD)
S4 App design and use
S5 Statistical analysis of *Drosophila melanogaster* Samarkand wings
S5 Statistical significance
S6 Code and data availability
S7 Supplementary figures

### S1 Deep learning based image segmentation

#### S1A. U-Net training details

MAPPER utilizes U-Net for training. U-Net (Falk et al. 2019) was trained using ground truth labels from 1000 raw wing images. The ground truth was generated using ILASTIK by first training a pixel classifier to identify the four pixel classes: vein, intervein, wing marginal hair, and background. The segmentation masks generated using ILASTIK were then used to train the U-Net. The output layer of a pre-trained U-Net that was earlier trained for biomedical image segmentation (Ronneberger, Fischer, and Brox 2015) was then modified to detect multiple classes. A cross entropy cost function was used (Goodfellow, Bengio, and Courville 2016). Training was carried out in PyTorch and the implementation was done on GPUs. The raw images used for the training were imaged using an Aperio (Leica) slide scanner at 5X resolution.

#### S1B. Deployment of a trained deep learning model on raw wing image data

The trained model can be found in the attached <u>GitHub repository</u>.

### S2 Training the intervein classifier

#### S2A. Generation of training data

MATLAB’s Image labeler app was used to generate ground truth labels for training an intervein classifier. Adult wing images were imported into the application, and the polygon tool was used to define each intervein regions. A total of ten different classes were created that included the seven intervein regions in a normal adult wing blade, along with the cases of cross vein defects where either the anterior or posterior cross vein of the wings are missing (Figure S4A). A class where the L_5_ vein fails to meet the margin of the wing blade, named as an L_5_ end defect, has also been included. A total of 81 wing images belonging to five different classes present in the “<u>Source/ InterveinClassification/GeometricFeatures/ TrainingData</u>“ were used for generating a total of 496 different objects belonging to the 10 intervein classes. The images used for labeling were marked and saved in “<u>Source/InterveinClassification/GeometricFeatures/TrainingData</u>”. The five different classes and their folder names are:

- NormalPixelLabelDat
- ACVPixelLabelData
- PCVPixelLabelData
- ACVPCVPixelLabelData
- ACVEnddefectPixelLabelData.

#### S2B. Feature extraction

Two different kinds of feature extraction methodologies were followed while extracting features from different intervein components within the wing. The first one comprises basic geometric features. These include normalized intervein areas, eccentricity and the aspect ratio of each intervein component. The feature vector for all the training data can be extracted by running “<u>Source/InterveinClassification/GeometricFeatures/InterveinClassifierGeo.m</u>”. The feature vector is stored as an array where each row is a sample, and the three different columns contain the features. The labels for these are stored in the array named “ResponseTraining”. Another methodology is when we used Elliptic Fourier Descriptor-based shape features (Kuhl and Giardina 1982). The number of features in this case depends on the number of harmonics to define an EFD model for the interveins. In our studies, we used a total of 15 EFD harmonics that resulted in a feature vector of size 496x(15×4).

#### S2C. Training a machine learning classifier

The features extracted from the training data are imported to the MATLAB’s classification learner app. All the models available were trained on the training data with a ten fold cross-validation. Based on the geometric features, a weighted KNN performed the best, while based on the EFD features a Fine KNN was seen to have the most accurate classification (Figure S4C, Figure S4D). Both the models were saved for future use in intervein classification. Details on how we further choose the best model for our intervein classification has been described in the main text.

### S3 Morphological feature extraction using the labelled interveins

#### S3A. Intervein geometric features

The machine learning-based intervein classification pipeline classifies and labels each intervein region. Then, MATLAB’s regionprops tool extracts geometric features of each intervein. Geometric features like area, aspect ratio and circularity circularity have been extracted and stored in the table containing wing features.

#### S3B. Trichome density of interveins

Trichomes are hairlike structures that are present in the *Drosophila* wing blade. The location where the base of the trichome originates is of darker intensity compared to the neighboring pixel. For each pixel present in the intervein region, we checked if it was a local minima in the defined window size containing the set of neighboring pixels. The window size was chosen as a circular patch of radius equalling half the length of a trichome. The same has been defined as a user input parameter as the length of hair in pixels varies with resolution of imaging. Once the location of trichomes is defined, we display the local trichome density using 2D histograms.

#### S3C. Landmark position-based features

##### S3C-I. Length of proximal-distal axis

The calculation of length of proximal-distal axis requires definition of veins L_3_ and L_4_. Vein L_3_ is reconstructed through morphological operations on I_2_, I_3_ and I_4_ in MATLAB. Definitions of the interveins can be found in Figure S1. Interveins I_3_ and I_4_ are joined first by using dilating the boundary pixels and then by eroding by the same amount the joined components to form intervein I_3,4_. In a similar way I_2_ and I_3,4_ are joined together to form I_2,3,4_. I_2_ and I_3,4_ are then subtracted from I_2,3,4_ to obtain the reconstruction of vein L_3_. The same has been shown in Figure S5. We use a similar methodology to define vein L_4_ as well. The veins are then skeletonized and the vein endpoints are then extracted by locating pixels in the skeleton having maximum distance between them. The oriental of the wing allows identification of the proximal and distal ends of these veins. The average position of the proximal ends and distal ends is then used to get the proximal-distal axis. Euclidean distance between the two points defines the length. (See Figure S5 for more details on how segmentation of veins are constructed).

##### S3C-II. Length of anterior-posterior axis

Specification of vein L_2_ and L_4_ is required to define the anterior-posterior axis of the wing blade. We follow a similar strategy as described above to define veins L_1_ (I_1_ and I_2_) and L_4_ (I_5_,I_6_, and I_7_). The distal ends of veins L_2_ and L_4_ are then used to define the anterior-posterior axis of the wing.

#### S3C-III. Distance between veins L_3_ and L_4_

Distal ends of veins L_3_ and L_4_ are used to estimate the distance between veins L_3_ and L_4_.

#### S3D. Elliptic fourier descriptors

EFDs have been used for a wide number of studies related to plant/leaf evolution and shape analysis of organs (Kuhl and Giardina 1982). We used Elliptic Fourier Descriptors (EFD) as an alternative for a robust translational and rotational invariant representation of wing shape. EFDs are found by fitting a fourier series to the periodic function obtained from the closed *Drosophila* wing peripheral contour. The coefficients of the Fourier series act as features as each of them carry a local shape property. The original EFD description is a scale, translation, and rotation invariant shape descriptor (Kuhl and Giardina 1982). We modified the original algorithm, so it is sensitive to size changes. Details about this implementation can be found in the design section of the main text. To find the appropriate number of terms for representing the closed contour of the *Drosophila* wing blade, we varied the number of harmonics and measured the errors between the EFD reconstruction of the wing and the actual boundary points (Figure S6). Boundary points have been extracted from the U-Net/ILASTIK generated segmentation mask. With an increase in the number of terms, due to overfitting, the error decreases and saturates to 0. A further manual inspection showed that 20 harmonics are sufficient for representing any wing blade. In summary, EFDs allow us to measure specific local changes within the wing corresponding to a genetic or a pharmacological perturbation.

### S4 App design and use

#### S4A. File formats and folder organization for running the app

For any individual processing or batch processing application, the raw wing images have to be saved in the form of a ‘.tiff’ format file. Alternatively, this can also be achieved in the wing processing/hinge removal step explained in the next section. Additionally, the segmentation masks have to be saved in a ‘.png’ format. MAPPER also requires unique nomenclature for the raw wing image and the generated segmentation mask (i.e. if a raw wing image is saved as “wing.tiff”, the corresponding segmentation mask has to be named “wing_Simple segmentation.png”. This kind of a nomenclature can be generated using ILASTIK by modifying the batch processing options. Processing wing images using U-Net, for generating the segmentation masks, already saves images in the correct format and naming convention.

#### S4B. Wing preprocessing

The adult *Drosophila* wing consists of a main wing blade area that mainly comprises of intervein and vein regions. The wing blade is connected to a hinge that joins the wing blade to the *Drosophila* body. We developed a methodology for every wing to be analyzed consistently by cropping out the hinge portion from the adult blade. This is done by selecting two landmark points that define the hinge endings in the anterior and posterior half of wing as shown in Figure S7. Using these two points, a polygon is drawn around the blade that encompasses the hinge region. This operation is carried out using a ‘roipoly’ function in MATLAB. A value of 0 is then assigned to the drawn polygon to label the hinge area. Unlike other existing tools, MAPPER does not require the images to be oriented in a particular fashion. The user needs to define the input path of the folder containing the raw uncropped wings for using this function which is embedded in the button “Remove wing hinge” shown in Figure 3D. User images should be placed within the “userData” subfolder to ensure proper functionality. The function also generates the output in the desired ‘.tiff’ format.

#### S4C. Input parameters

IP - Individual processing, BP - Batch processing. The wing process abbreviations IP and BP indicates if the parameters need to be defined while running them. Default parameters have been set for images taken at a 4x objective lens, but can be adjusted for images taken at other objectives.

##### i) Size of wing filter (IP, BP)

The intervein and vein regions are separated during the image segmentation process. The pipeline merges the two label definitions to generate a segmentation mask defining the complete wing blade. The joining of labels is followed by morphological operations in MATLAB to smooth out and filter the segmentation mask of the wing blade. More specifically, erosion and dilation are performed to smooth out the wing periphery. The parameter used in the app defines the radius of a disc shaped structuring element used for this particular morphological operation. A filter size of 8 has been seen to perform well for the images taken at 4x resolution. A typical value of this filter could be chosen as the thickness of veins (in pixels).

##### ii) Size of intervein filters (IP, BP)

The parameter used in the app defines the radius of a disc shaped structuring element used for erosion and dilation operations to smooth out the intervein boundaries. A filter size of 5 has been seen to perform well for the images taken at 4x resolution. A typical value of this filter could be chosen as half the thickness of veins (in pixels). The thickness of the vein can be approximated using FIJI/ImageJ.

##### iii) Threshold trichome distance (IP, BP)

Trichomes are hairlike structures patterned across the interveins. Their relatively darker appearance relative to the intervein region allows it to be distinguished as a minima in the image intensity plane. A local thresholding operation is used to count the number of minimas, and which gives the number of trichomes in the wing. Additionally, to avoid double counting we refine the list of minimas based on the distance between the neighbouring hairs. If the distance between the two local minimas is less than typical size of a trichome, it leads to deletion of one of the identified trichomes from the list. The parameter is named threshold trichome distance, and a value of 2 is seen to work best on wings imaged at a resolution of 4x.

##### iv) Number of harmonics (BP)

The parameter is used to define the number of harmonics or number of terms in the EFD definition of the wing blade. 20 harmonics was sufficient to correctly define the wing blade morphology (Figure S3).

##### v) Conversion factor (IP, BP)

The parameter is used to define the number of pixels in one unit of measurement. It is used to give output of quantities such as area and length in the user defined length unit of length. A default conversion factor of 2 microns per pixel is set in the tool.

##### vi) Label name (IP)

The text input requires user-based naming of an image that will be saved later. The image printed out consists of the labelled intervein regions and their corresponding masks. Users also need to define the file extension along with the file name (i.e. ‘Sample_LableWing.png’).

### Individual Wing Processing (Figure 3B)

#### S4D. User operations

##### i) Load Raw Image

The button directs the user to select the single raw wing image they want to process. The image loaded can be viewed in the “Raw Image” titled figure panel in the app.

##### ii) Load Segmentation Mask

The button directs the user to select the corresponding segmentation mask for the raw image they want to process.

##### iii) Classify interveins

The button executes the support vector machine (SVM)-based, pre-trained intervein classification scheme described in the main text. The results can be displayed in the “Labelled Wing Image” figure panel of the app. The area and number of trichomes present in each classified intervein, along with the intervein id, is printed in the table below the “Raw Wing Image” figure panel. Overall wing blade area and average trichome density of the intervein regions are also given as an output of this operation. The output is displayed in a text panel right below the user operations panel.

##### iv) Full Process

The button directs the user to select a single raw image for processing. Upon selection, the segmentation mask will be loaded, and the interveins will be classified automatically. The naming convention of the segmentation mask must be consistent with the raw image file name, as detailed in section S4A. Should the segmentation mask be labeled differently, the button “Load Segmentation Mask” should be used.

#### S4E. Biological axes

##### i) AP axis

The button directs the pipeline to calculate the length of the anterior-posterior (AP) axis of the *Drosophila* wing. The numerical output can be displayed in the output panel located just below the buttons. The AP axis is also overlaid on the raw image, and can be seen in the figure panel titled “Biological axes display”.

##### ii) PD axis

The button directs the pipeline to calculate the length of the proximal-distal (PD) axis of the *Drosophila* wing. The numerical output can be displayed in the output panel located just below the buttons. For visualizing the location of the PD axis, it is also drawn over the raw wing image. The resulting raw image with annotated PD axis can be seen in the figure panel titled “Biological axes display”.

##### iii) d(L_3_-L_4_)

The button directs the pipeline to calculate the distance between L_3_ and L_4_ veins at the distal most end of the *Drosophila* wing. The numerical output can be displayed in the output panel located just below the buttons. The length of the axis is also overlaid on the raw image, and can be seen in the figure panel titled “Biological axes display”.

### Batch processing (Figure 3C)

#### S4F. User operation

##### Batch processing

The button is used for batch processing of images sets. The folder containing the wings and corresponding segmentation masks have to be stored in their own folder placed inside the “userData” subfolder beforehand. When the button is pressed, the user is directed to select the folder containing the wing images. It is best to store images of wings belonging to different genotypes/populations in separate folders. A table containing all the features (description in section SI3) is saved in a .mat file labeled “T.mat”, and a .csv file labeled “Wing_Measurements_Batch_Output.csv”. Further, the EFD coefficients are saved in a .mat file labeled “EFDBatch.mat”, and a .csv file labeled “EFD_Batch_Coefficients.csv”. An image showing labelled intervein regions for each image, is also saved in the directory that contains the selected wing images and segmentation masks. This is to allow users to ensure quality control of proper segmentation and classification. For the calculations described below, an assumption that wings located in the same subfolder belong to the same genotype/population is used.

#### S4G. Average area figure panel

Average area of wings contained in each subfolder analyzed is plotted as bar graphs and displayed in this figure panel in the batch processing section of the app.

#### S4H. Hits z-score

Geometric features (see feature extraction in section SI3) of the intervein regions are used to define a feature vector for each wing. The features of each wing in a specific folder are averaged out to define the average feature vector for a specific genotype/population. Mahalanobis distance is then calculated between the average feature vector of a subfolder and the average features of all the wings analyzed to define the z-score. The genotypes/populations with maximum z scores are shown in the text panel “Hits z-score”.

### S5 Statistical Analysis of Samarkand strain *Drosophila* wings

We processed 128 adult wing images of *Drosophila melanogaster* from the Samarkand strain (Figure 2, Figure S10) (Sonnenschein et al. 2015)(Sonnenschein et al. 2015)(Sonnenschein et al. 2015). In this validation test, MAPPER was used to highlight the shape and size differences between the male and female populations. Geometric features and EFD features were separately analyzed to highlight the application of EFD in estimating local shape changes in wing. Principal Component Analysis (PCA) (Wold, Esbensen, and Geladi 1987) carried out on the geometric features revealed that the maximum variance within data was distributed majorly between the first two principal components (89.4%) (Figure 2E, Figure S10C-D). PCA also revealed that the total area of the wing and total trichome density had maximum loading towards Principal Component 1 (PC1) with total wing area having a negative PC1 loading towards the female population, and total trichome density having a positive PC1 loading towards the male population (Figure S10C-F). Interestingly, the distance between longitudinal veins L_3_ and L_4_, d(L_3_-L_4_), did not correlate with other wing features for either population (Figure S10A). Further, d(L_3_-L_4_) did not have positive or negative loading for PC1, which explained most of the variance in the data (Figure S10B-D). This is contrary to the other biological axes measurements for the AP and PD lengths that had negative PC1 loading towards the female population. The differences between males and females were further characterized by plotting the known result that wings of a *Drosophila* female are greater in size than the male wing (Figure 2F, Figure S10F). PCA analysis and comparison of the standardized total wing areas confirmed this (Figure S10C,D,F, *p*-value < 0.001). Conversely, males had a larger total standardized trichome density (Figure S10G, *p*-value < 0.001) that is revealed by direct comparison and PCA analysis (Figure S10C-D). Taken together, this suggests that high dimensional data provided by MAPPER can reveal unique features that are disparate between populations using PCA analysis. Clustering carried out using the first two principal components revealed the presence of two distinct clusters representing the male and female populations (Figure 2E). Further, performing t-distributed stochastic neighbor embedding (t-SNE) analysis revealed similar clustering distinctions between male and female populations (Figure S10E). High dimensional analyses, such as PCA and tSNE, allow for a systematic screening of phenotypic changes between two different populations, and can even be extended to multiple populations.

A similar approach was followed in analysis of the EFD-based features. PCA was applied on the EFD coefficients extracted for each wing. In this case, a total variance of about 97% was captured in PC1 alone (Figure 2G). This indicates that the variation was mainly distributed in one direction of linear combinations of the EFD coefficients. Thus, the high dimensional output of MAPPER coupled with PCA analysis can reveal wing shape features that are distinct between two populations. To investigate the importance of PC1 on wing geometry, we implemented reverse PCA on the principal components. Reverse PCA is carried out by adding the mean vector of the features to the matrix product of PCA projections (scores) and the transpose of the eigenvectors (Equation 1). This process enables mapping of the influence of principal components back to the original data.

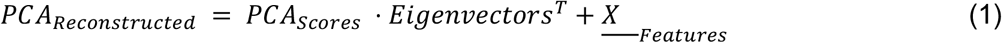

In our analysis, the first eleven principal components were used to reconstruct the EFD coefficients via reverse PCA. Since we are interested in understanding the importance of PC1, which explained most of the variance in the data, the standard deviation along this PC was calculated. Mean scores of the first 11 PC’s were then used to reconstruct a representative wing for the male and female populations (Figure 2I). 1.5 times the calculated standard deviation along PC1 was then added and subtracted to the representative wing to observe the effect of PC1 on wing shape. Reverse PCA was then used to approximate the EFD coefficients with varying PC1. Reverse EFD was then carried out to estimate the contour of the wing blade. The reconstructed contours highlight that the major differences between the wings is mainly because of overall change in the wing blade area (Figure 2H). However, the reconstructed contour can be used as the mean representation of the samples. This approach is useful when analyzing subtle changes in shapes resulting from dysregulations in genetic pathways. These subtle differences are further characterized by clustering using Gaussian Mixture Models (Yang, Lai, and Lin 2012) (GMM) where the presence of two distinct clusters is revealed (Figure 2G). Both of these high dimensional plots reveal that there are unique and identifiable distinctions between male and female populations in regard to wing shape.

### S6 Tests for measuring statistical significance

One way analysis of variance (ANOVA) was first used to test the hypothesis if the means of the groups compared were equal or not (Fisher 1992). However, ANOVA alone cannot be used to comment on statistical significance of comparisons between any two subgroups. Therefore, we further used a multiple group comparison test for the statistics generated by ANOVA. Tukey’s honestly significant difference procedure was used for this task to identify differences in means among all subgroups (Tukey 1949). Bonferoni-Holm correction was then applied on the generated *p*-values from the previous test in order to adjust for Type I error in statistical testing (Holm 1979). All the steps were carried out using MATLAB. Additionally, a Bartlett test was used to compare the variances between any of populations.,

## Supplementary Figures

**Figure S1.**
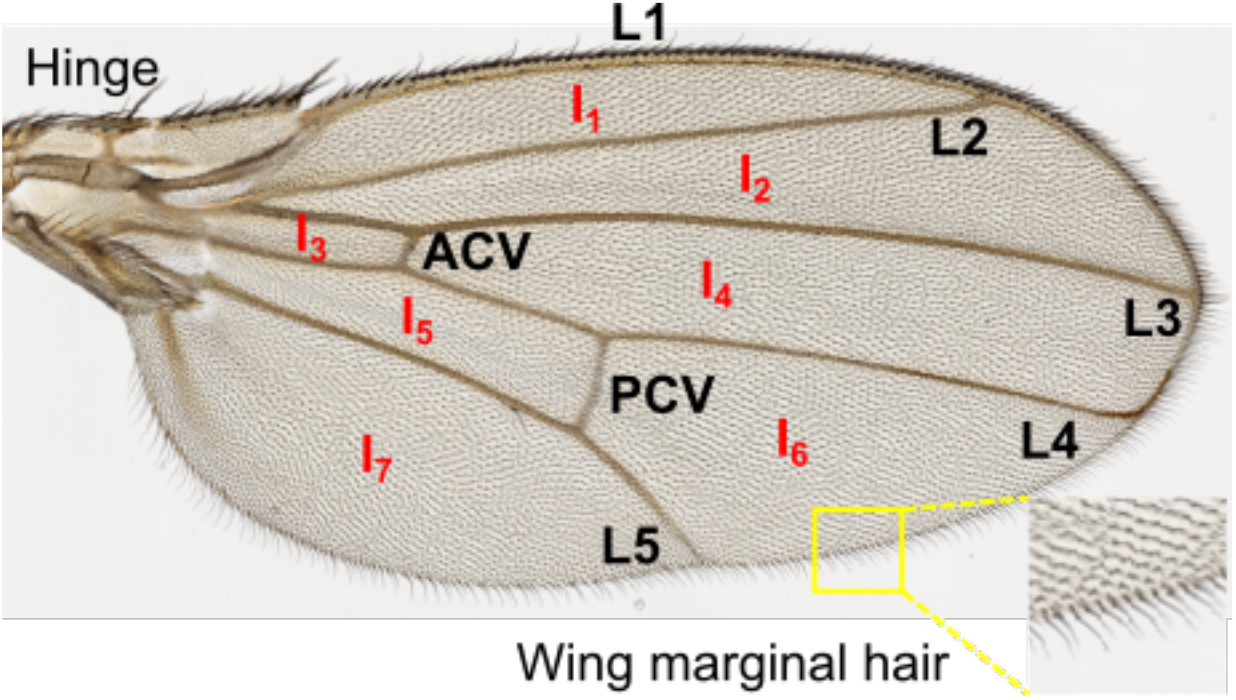
Components of adult *Drosophila* wing blade. The adult *Drosophila* wildtype wing has distinct morphological features: longitudinal veins L_1_-L_5_, anterior cross vein (ACV), and the posterior cross vein (PCV), that we rely on to study the effect of specific genetic perturbation on morphogenesis. Veins also enclose seven specific intervein regions (I_1_ - I_7_). The periphery of the wing is surrounded by hairs. The intervein regions and veins are also patterned with small hair like structures, called trichomes. The wing is attached to the fly body through a hinge.

**Figure S2.**
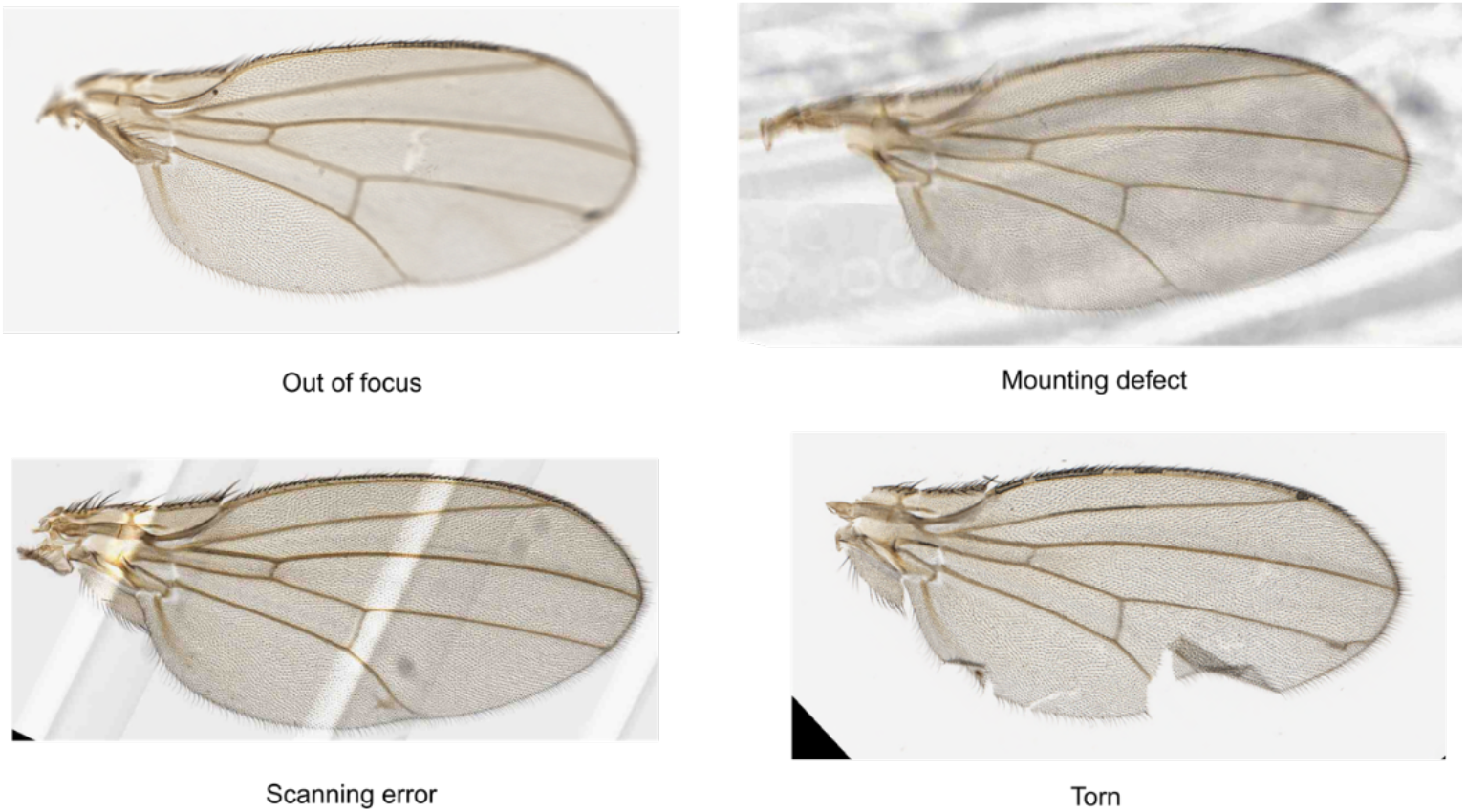
Defective wing images that are excluded from the pipeline. Shown here are examples of wing images that contain various image artifacts or sample preparation defects. The MAPPER pipeline is able to recognize these defects and remove the corresponding images from the analysis.

**Figure S3.**
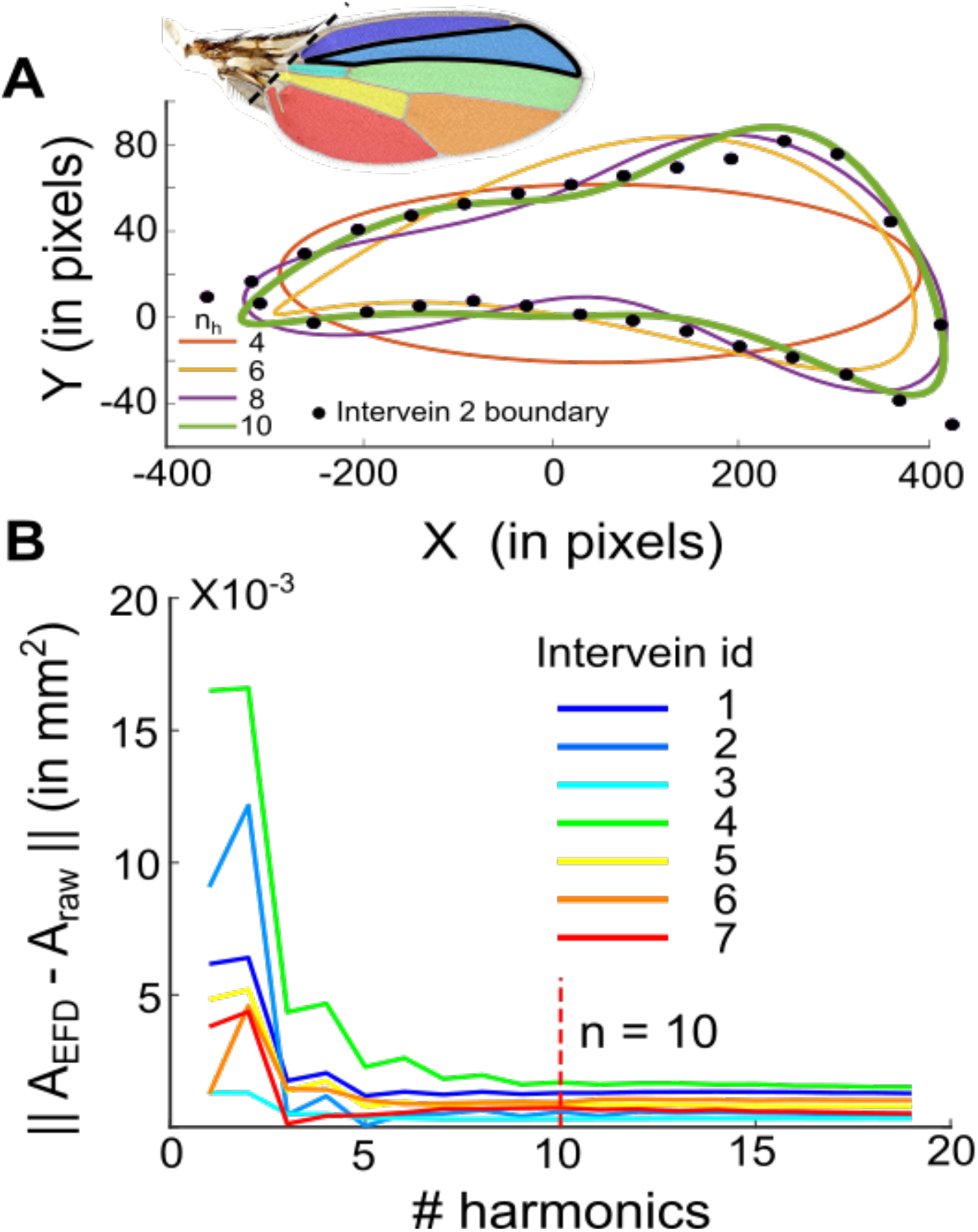
A total of ten terms in EFD description is sufficient to define the geometry of an intervein. **(A)** Variation of number of terms in the elliptic fourier descriptors (EFDs) on accuracy of fit for specifying the intervein enclosed between veins L_2_ and L_3_ (I_2_). As the number of terms increases, the EFD fit approaches the true intervein shape. **(B)** Variation in error with an increase in number of terms in the EFD description for each intervein. The error is approximated by calculating the absolute difference between area enclosed by the EFD fit and actual intervein area.

**Figure S4.**
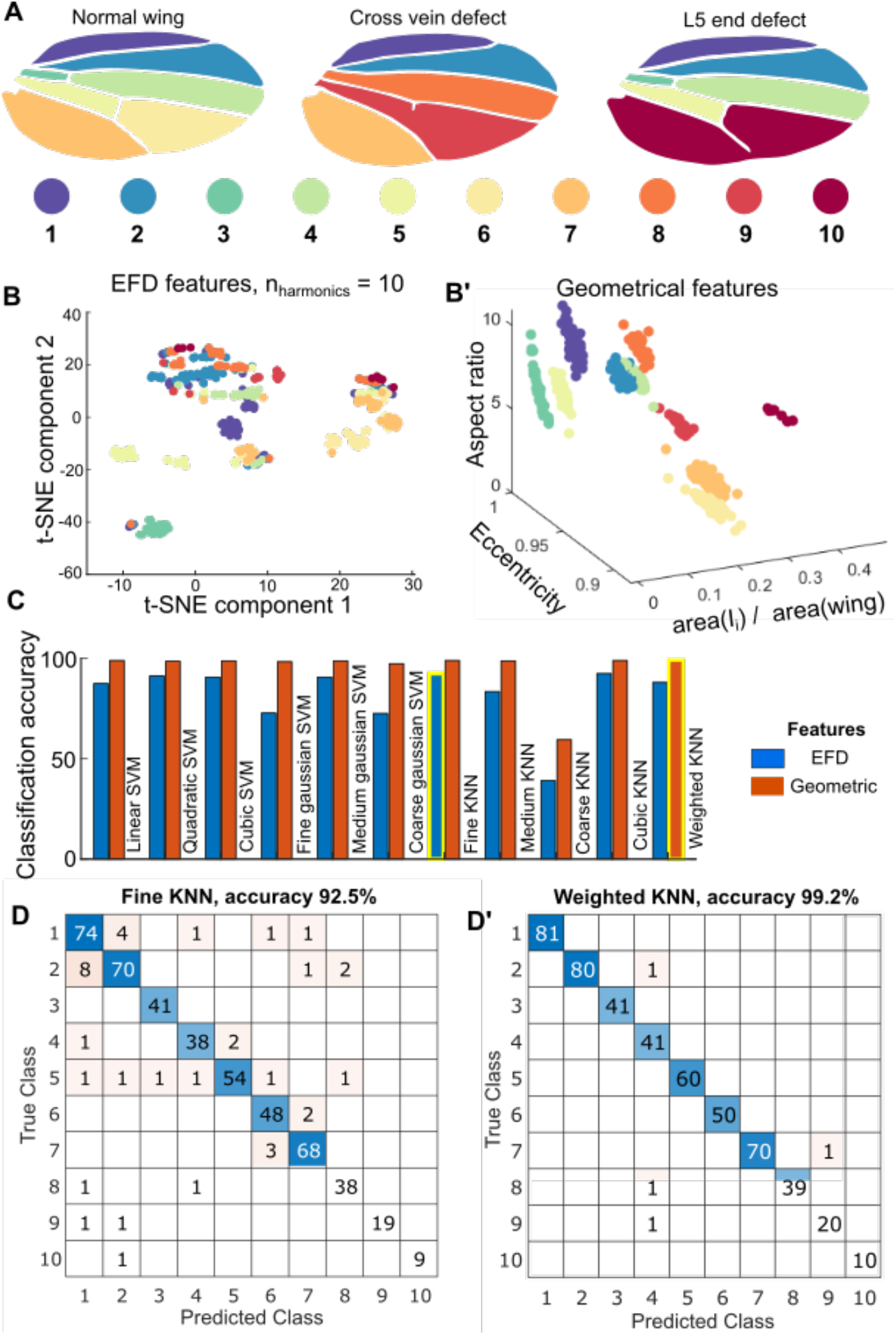
KNN trained on geometric features performs best for intervein classification. **(A)** Definition of each intervein region used while creating the training data for the purpose of intervein classification. Vein defects led to additional intervein classification regions. **(B)** t-SNE representation of EFD features extracted from each individual intervein region from a batch of wing images used for generating the training dataset. Clear clustering is present in the different intervein regions. **(B’)** Geometric features such as normalized intervein area, eccentricity, and aspect ratio are also used for training an alternate intervein classifier. **(C)** Performance of different support vector machines (SVMs) and k-nearest neighbor algorithms (KNNs) present in MATLABs classification learner application based on the two different sets of features used for the purpose of training. Weighted KNN performed the best compared to other methods. **(D-D’)** Confusion matrices for best models based on the two different sets of features used. (D) uses the EFD-based features while (D’) uses the geometrical features of individual intervein.

**Figure S5.**
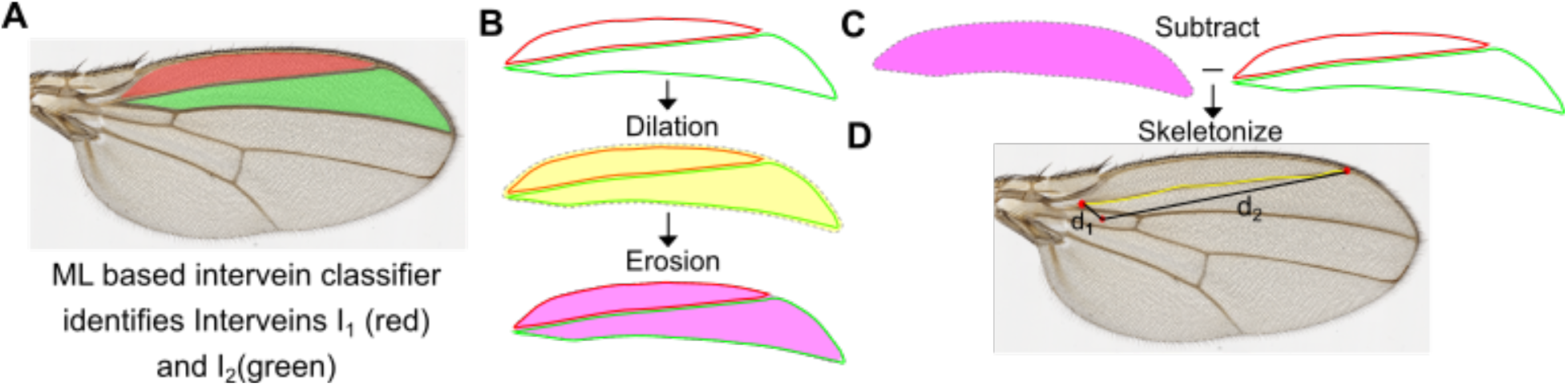
Methodology for extracting proximal and distal ends of longitudinal vein L_2_. **(A)** The intervein classification pipeline identifies intervein region I_1_ and I_2_. **(B)** Binary masks for intervein regions I_1_ and I_2_ are dilated in order to fuse them together. A disc structuring element with radius equaling vein thickness was used for this. The dilated image is then eroded by the same structuring element. **(C)** The binary masks for I_1_ and I_2_ are then subtracted from the binary mask obtained after dilation and erosion, and the resulting mask is then skeletonized to obtain the definition of vein L_2_. The distance of the ends of the vein are calculated from the center of intervein region I_3_ to estimate the proximal and distal ends of vein (see Figure S1A).

**Figure S6.**
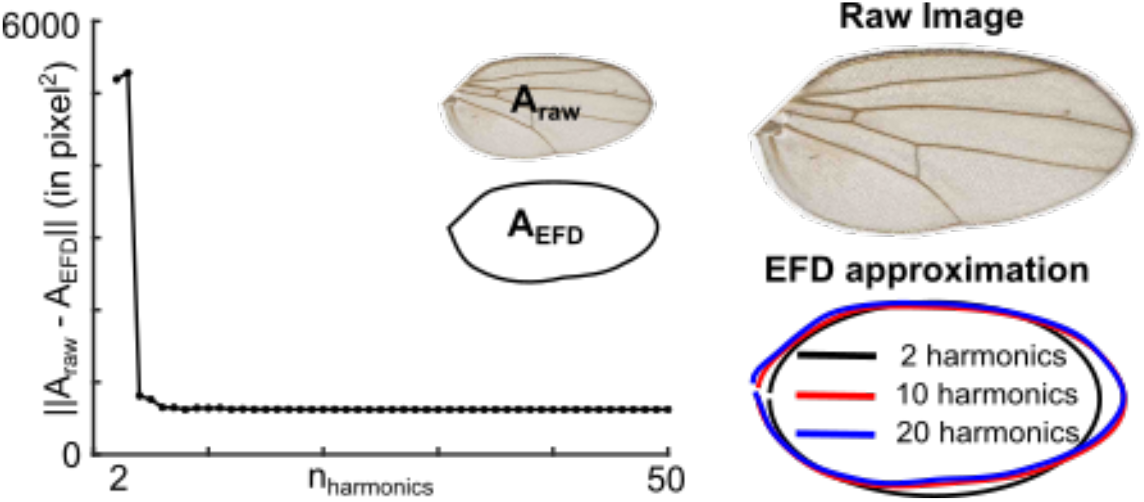
EFD description of the global wing shape. Elliptic Fourier Descriptor (EFD) coefficients are determined to fit the perimeter of the adult wing blade. Left-hand side graph shows the variation of error with an increase in the number of harmonics or the terms used in EFD approximation. Error is approximated by calculating the absolute differences between the actual area of the wing and the area enclosed by an EFD fit. Right hand side panel shows the actual representation of the effect of increasing the number of harmonics on overall fit.

**Figure S7.**
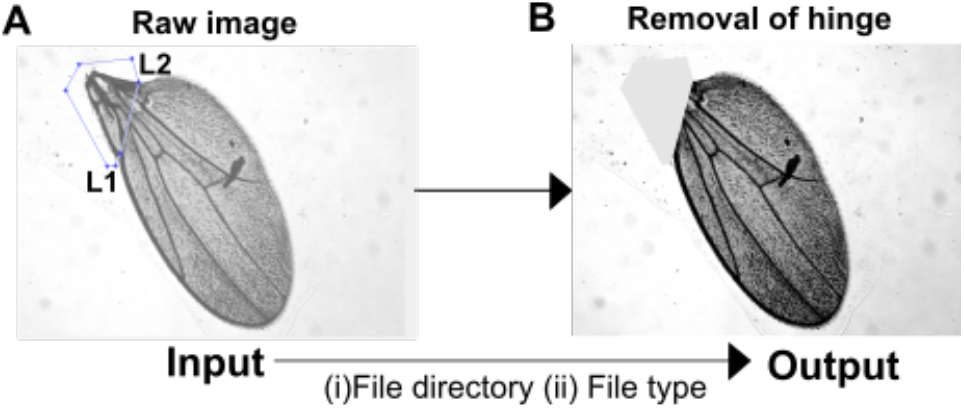
Wing hinge removal. **(A)** Landmark points L_1_ and L_2_ marking the anterior and posterior ends of the hinge region are used to draw a polygon using MAPPER’s wing hinge removal button (see section SI4B). **(B)** The polygon is then cropped out of the raw wing image for full processing by MAPPER.

**Figure S8.**
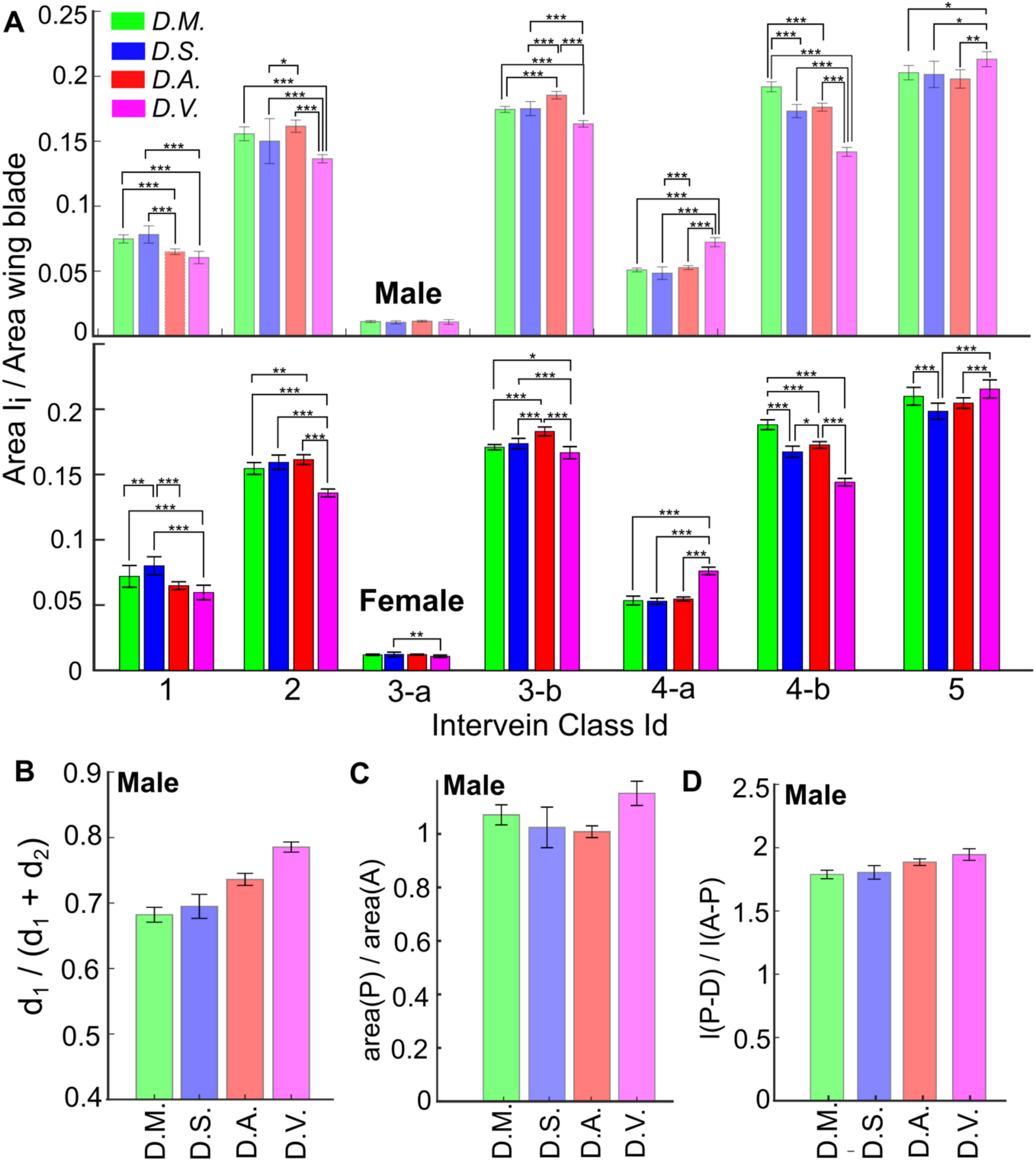
Differences in intervein areas and landmark positions in different *Drosophila* species identified by MAPPER. **(A)** Ratio of individual intervein areas normalized to the overall wing blade area. Each subgroup of the bar plot indicates the quantification of the ratio for four different species indicated in the legends. The analysis has been carried out for the seven distinct intervein regions, whose labels are indicated in the x-axis label at the bottom of the panel. Top and bottom rows indicate analysis done for the male and the female sexes of each individual species respectively. **(B)** Quantification of shift in posterior cross vein position in male wings (L_1_ is defined as the segment of L_5_ from the proximal to posterior cross vein, L_2_ is defined as the segment of L_5_ from the posterior cross vein to the end of L_5_). **(C)** Relative anterior to posterior areas in female wings for each species. **(D)** The ratio of the length of the proximal-distal axis and the anterior-posterior axis for females from different species. Error bars are representative of standard deviation. (*** *p*-value < 0.001, ** *p*-value < 0.01, * *p*-value < 0.05)

**Figure S9.**
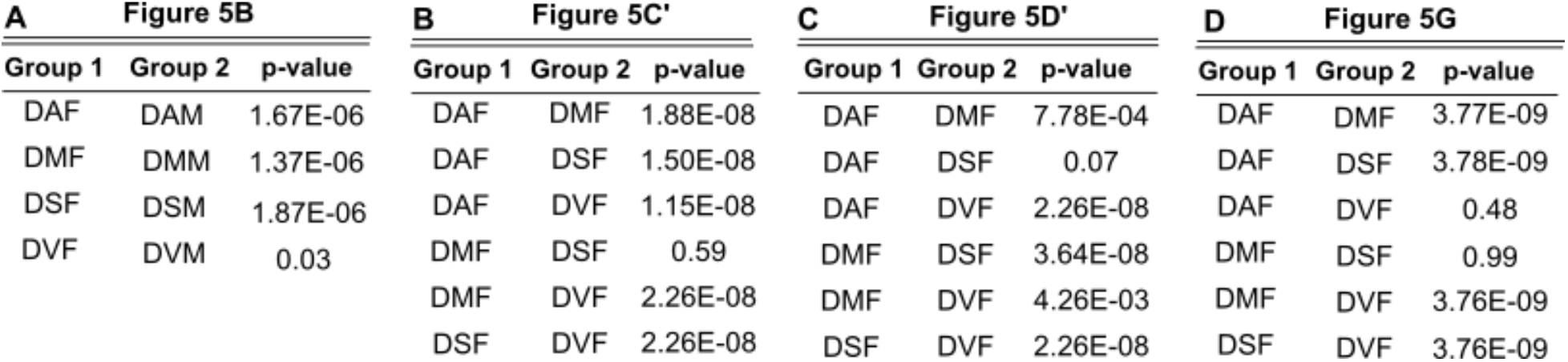
Statistical testing for differences in wing features of different *Drosophila* species. Tables containing *p*-values from a Tukey honestly significant difference statistical test performed to check statistical significance between wing features of different *Drosophila* species. The heading of the plots indicate the main figure number the table is meant for (Figure 5). Further details about the statistical tests can be found in section SI5.

**Figure S10.**
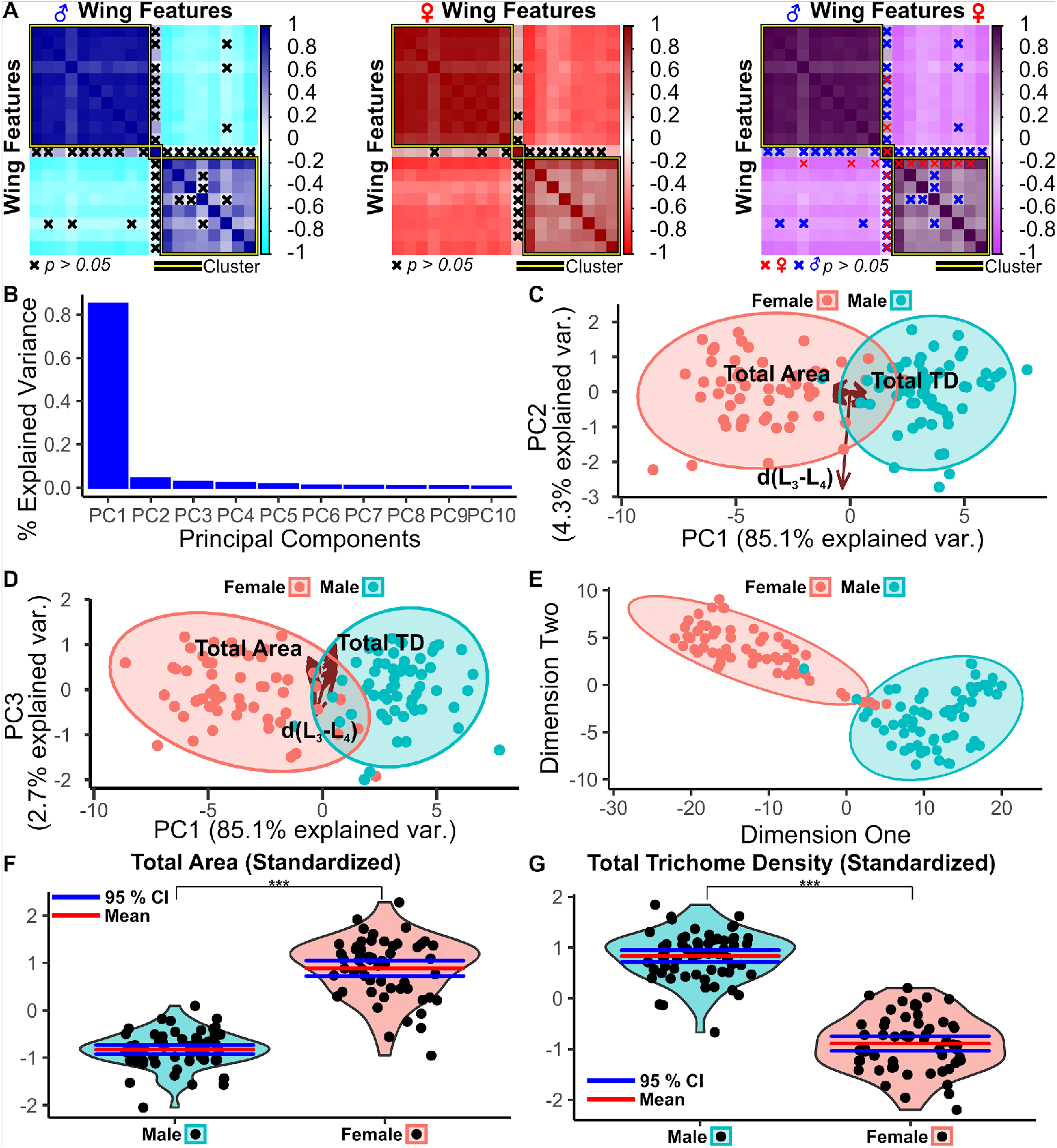
MAPPER provides high dimensional analysis capabilities to identify unique morphometric features. *Drosophila* adult wings from Samarkand strain males and females were analyzed using MAPPER’s batch processing capabilities (Sonnenschein et al. 2015). **(A)** Correlation plot of male, female, and averaged male/female morphometric features respectively. Insignificant correlations are denoted by “x” marks. Hierarchical clustering revealed three clusters: wing areas, distance from the L_3_ vein to L_4_ vein d(L_3_-L_4_), and trichome densities. Males and females have similar clustering and correlations of features. **(B, C, D)** Principal component analysis (PCA) on centered and scaled wing features revealed most of the variance in the data is explained by PC1, with distinct clustering across PC1 between males and females. Arrows represent projection of eigenvectors for each feature, and ellipses are 95% confidence intervals (CI) of the data populations. **(E)** t-SNE analysis on wing features revealed distinct clustering between males and females. **(F, G)** Violin plots of standardized wing features identified in PC analysis. means (red) and 95% CIs (blue) means (red) are overlaid for comparison. (*** *p*-value < 0.001 via Mann-Whitney U Test)

**Figure S11.**
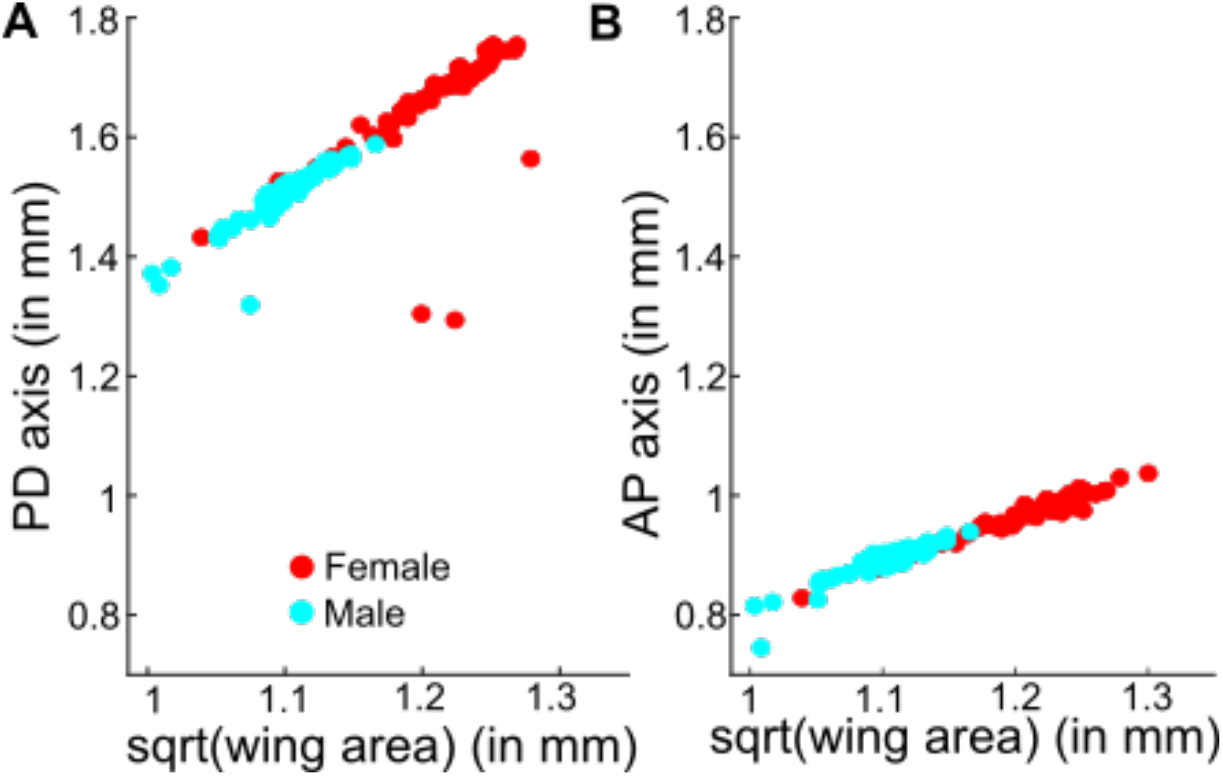
AP axis and PD axes scale uniformly with respect to area of the wing. Analysis was carried out for a population of male and female wings belonging to *Drosophila melanogaster* Samarkand strain. The points are color coded according to sex where red indicates fremale and blue the male population of wings. **(A, B)** Variation of proximal-distal axis and anterior-posterior axis with respect to square root of area of wing blade.

## Notes

https://mselab.github.io/MAPPER_Quantitative/

### Competing Interest Statement

The authors have declared no competing interest.

